# Dynamic transient brain states in preschoolers mirror parental report of behavior and emotion regulation

**DOI:** 10.1101/2024.01.22.576699

**Authors:** Lisa Toffoli, Natalia Zdorovtsova, Gabriela Epihova, Gian Marco Duma, Fiorella Del Popolo Cristaldi, Massimiliano Pastore, Duncan E. Astle, Giovanni Mento

## Abstract

The temporal dynamics of resting-state networks (RSNs) may represent an intrinsic functional repertoire supporting cognitive control performance across the lifespan (Kupis et al., 2021). However, little is known about brain dynamics during the preschool period, which is a sensitive time window for cognitive control development. The fast timescale of synchronization and switching characterizing cortical network functional organization gives rise to quasi-stable patterns (i.e., brain states) that recur over time. These can be inferred at the whole-brain level using Hidden Markov Models (HMMs), an unsupervised machine learning technique that allows the identification of rapid oscillatory patterns at the macro-scale of cortical networks (Vidaurre et al., 2018). The present study used a HMM technique to investigate dynamic neural reconfigurations and their associations with behavioral (i.e., parental questionnaires) and cognitive (i.e., neuropsychological tests) measures in typically developing preschoolers (4-6 years old). We used high density EEG to better capture the fast reconfiguration patterns of the HMM-derived metrics (i.e., switching rates, entropy rates, transition probabilities and fractional occupancies). Our results revealed that the HMM-derived metrics were reliable indices of individual neural variability and differed between boys and girls. However, only brain state transition patterns toward prefrontal and default-mode brain states, predicted differences on parental-report questionnaire scores. Overall, these findings support the importance of resting-state brain dynamics as functional scaffolds for behavior and cognition. Brain state transitions may be crucial markers of individual differences in cognitive control development in preschoolers.

**Keypoints:**

- HMM-derived metrics are reliable hallmarks of individual neural variability and show gender-related differences.
- Brain state transition patterns toward prefrontal and default-mode brain states predict differences on parental-report questionnaires scores.
- Brain state transitions may be crucial markers of individual differences in cognitive control development in preschoolers.

## Introduction

Resting-state brain activity can be defined as an intrinsic functional repertoire of spontaneously active fluctuations with specific spatiotemporal patterns (Baker et al., 2014). These endogenous dynamics provide the functional substrate for exogenous and/or task-evoked activations and are related to behavioral and cognitive outcomes (Avery et al., 2020; Barnes et al., 2016; Cai et al., 2018; Jones et al., 2022; Poole et al., 2016). The brain’s functional organization has largely been investigated across the lifespan using a variety of neuroimaging techniques, such as fMRI (Smith et al., 2013; Smitha et al., 2017), MEG (Brookes et al., 2011; Wens et al., 2014) and EEG (Duma et al., 2021, 2022; Yuan et al., 2016). These studies have yielded novel insights highlighting a diffuse-to-local developmental shift in activation patterns accompanied by the strengthening of long-range connectivity at the expense of local connectivity (Fair et al., 2007; Uddin et al., 2010). Crucially, the capability of networks to dynamically modulate their functional organization stands as a scaffolding property, enabling adaptive responses to environmental demands through goal-directed forms of behavior (Bassett et al., 2006; Braun et al., 2015). Put simply, if functional networks are to be a meaningful scaffold for cognition then they must have the capacity to organise and re-organise at a sub-second timescale. This is particularly relevant for dynamic and flexible processes subsumed under the banner of cognitive control (CC; Diamond, 2013; Braem & Egner, 2018). However, the relationship between resting state network (RSN) reconfiguration and CC remains unclear. Many studies suggest that CC relies on the flexible interplay between the cognitive control network (CCN) - a distributed circuit of regions (including fronto-parietal areas) - and the default-mode network (DMN) (Alvarez & Emory, 2006; Dwyer et al., 2014; Mansouri et al., 2017; Niendam et al., 2012; Rottschy et al., 2012; Satterthwaite et al., 2013). The CCN activates in task-positive conditions (i.e., when participants are actively engaged in a task), when cognitive control is required to achieve a goal or resolve a conflict, whereas the DMN activates in task-negative, resting-state conditions and has been associated with self-referential processing and states of mind-wandering (Raichle, 2015). The progressive segregation between these two networks across development supports cognitive and behavioral refinement over time (Breukelaar et al., 2020). As development progresses, the energetic costs associated with transitioning toward CCN regions decrease, possibly due to structural white matter maturation (Cui et al., 2020; Tang et al., 2017).

As it has been shown that the interplay between CCN and DMN is crucial for task-related CC, their intrinsic dynamic interaction as measured during resting state may be used as a proxy for understanding CC efficiency across the lifespan (Hutchison & Morton, 2016; Kupis et al., 2021; Nomi et al., 2017). Cortical networks are characterized by a fast timescale of synchronization and switching, resulting in quasi-stable brain state patterns that recur over time (Allen et al., 2014; Liu & Duyn, 2013), which provides fundamental scaffolding for cognitive processes (Brookes et al., 2014; Hutchison et al., 2013; Bressler & Tognoli, 2006). However, the majority of studies inspecting these phenomena have used methods (e.g., sliding window approaches, clustering techniques) that do not track millisecond-scale temporal dynamics of latent brain processes (Taghia et al., 2018). Methods that are able to capture finer timescales may be necessary to investigate the dynamic nature of activity across the whole brain. Individual differences in neural dynamics might be particularly relevant during early childhood, which is a sensitive window for CC development (Diamond, 2013). During this age period, improved inhibition, flexibility and working memory are supported by functional changes in CCNs, but little is known about their underlying fine-grained dynamics. A promising computational approach that is able to track rapid oscillations within cortical networks is Hidden Markov Modelling (HMM), an unsupervised machine learning technique that allows the identification of mutually-exclusive discrete patterns of whole-brain spontaneous activity that recur over time (Vidaurre et al., 2018). Importantly, HMMs allow the characterization of whole-brain transitions between discrete states of stable and coordinated activity. HMMs have been used to infer resting-state and task-related dynamic properties from a range of different neuroimaging techniques, such as fMRI (Dang et al., 2017; Goucher-Lambert & McComb, 2019; Hussain et al., 2023), MEG (Baker et al., 2014; Vidaurre et al., 2018; Quinn et al., 2018; Hawkins et al., 2020) and EEG (Obermaier et al., 2001; Williams et al., 2018; Dash et al., 2020; Marzetti, 2023). Previous studies have revealed that the dynamic properties of RSNs are related to a variety of cognitive and behavioral outcomes (Cabral et al., 2017; Taghia et al., 2018; Vidaurre et al., 2017). Interestingly, neural dynamics may be related to individual differences across neurodevelopment, including neurodevelopmental conditions like attention-deficit hyperactivity disorder (ADHD) and autism (Dammu & Bapi, 2019; Maya Piedrahita, 2021; Scofield et al., 2019). However, the majority of studies that have inferred HMMs from neuroimaging data have only included adult participants, and there has been a relatively limited focus on childhood development. A recent study used HMMs to investigate spontaneous neural dynamics in a transdiagnostic sample of 8-13 year old children (Zdorovtsova et al., 2023) and revealed a positive association between the complexity of individuals’ transitions (i.e., entropy) between RSNs and general cognitive ability. Additionally, the specific pattern of between-state transitions and time spent in each state further explained individual variations in cognitive ability: transitioning into or spending time within DMN-like and fronto-parietal/sensory states was associated with decreased and increased cognitive ability, respectively. Examining the fine-grained dynamics of RSNs at different developmental timepoints may yield further insights into how the relationships between neural dynamics, cognition, and behaviour change throughout early life.

Given the importance of the preschool-age period for CC development, the present study investigated the relationships between neural dynamics and measures of cognition and behavior in a sample of preschool-aged children (4-6 years). We used high density EEG (hdEEG) to better capture fast reconfiguration patterns of HMM-derived metrics. These included switching rates (i.e., rate of switching between brain states), entropy rates (i.e., complexity of individuals’ transitions), transition probabilities (i.e., the probability of switching from each brain state to all the others) and fractional occupancies (i.e., the overall proportion of time spent in each state).

As a first step we checked the within-subject reliability of our HMM-derived metrics. This holds significant importance in assessing the validity of these metrics. For this purpose, we recorded hdEEG during two separate resting-state sessions per subject, from which we derived HMM-based indices. Moreover, we assessed the effect of age and gender on these indices as prior studies suggested that both these factors may impact brain dynamics in older populations (Kupis et al., 2021; Scofield et al., 2019). Then, as the main goal we investigated the relationship between HMM-derived indices with broader measures of behavior (parental questionnaires) and cognition (cognitive tests). Again, previous studies present divergent findings; some indicate positive associations (Taghia et al., 2018; Zdorovtsova et al., 2023) while others report negative relationships (Cabral et al., 2017) between switching rates and cognitive performance. We expected positive relationships between CCN-like state transitions, fractional occupancies and cognitive ability, and negative associations between DMN-like state indices and cognitive ability (Zdorovtsova et al., 2023).

## Methods

### Participants

Fifty-nine participants were initially enrolled. Children (4-6 years old) were recruited from a local kindergarten in the Venetian Region of Italy. Non-verbal reasoning was assessed using the Coloured Progressive Matrices (CPM; Raven & Court, 1938) and children with a CPM score 2 or more standard deviations below the population mean were excluded (N = 1). Moreover, we excluded children for whom at least one of the two at-rest hdEEG recording sessions was discarded due to excessive noise/movement artifacts (N = 19). Participants had no known or diagnosed sensory, neurological or neuropsychiatric disorders and normal or corrected-to-normal vision. The final sample included 39 children (18 girls; 4-6 years, M = 4.8; sd = 0.7). The demographic characteristics of the sample are described in Figure 1.

**Figure 1.**
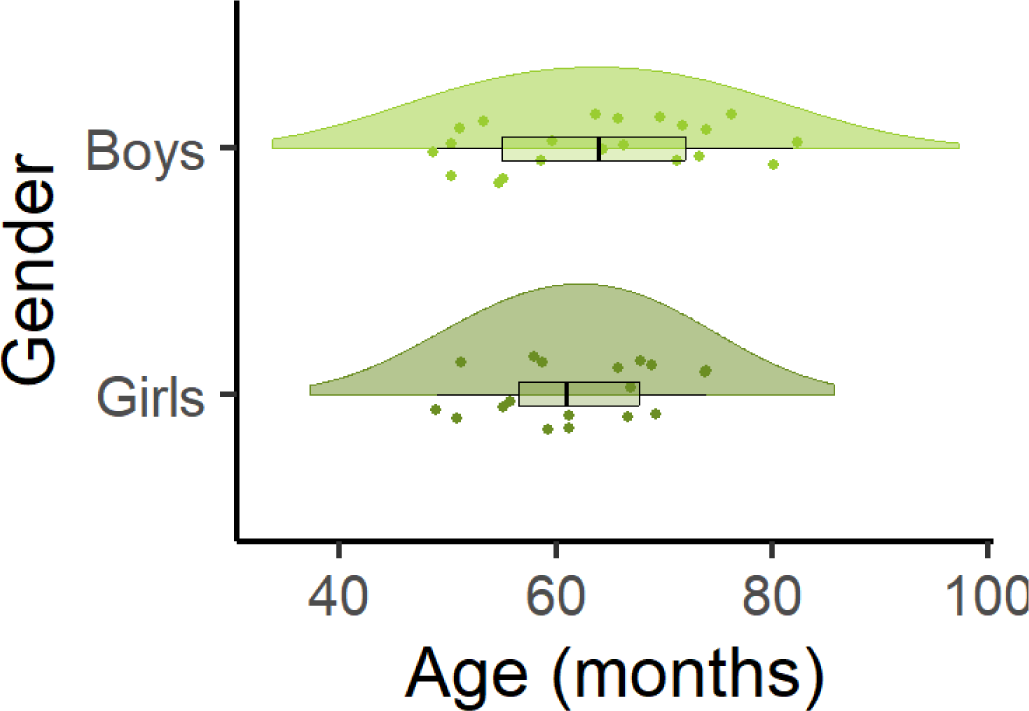
Raincloud plot depicting the distribution of age (months) across boys (light green) and girls (dark green). The box plots represent the interquartile range (IQR) with the median marked by a vertical bold line. The ‘cloud’ portions display the probability density of the data, providing insights into its distribution. Each data point is also shown to highlight individual observations.

### Ethics statement

Children’s parents provided written consent for their children’s participation. All experimental procedures were approved by the Ethics Committee of the School of Psychology of the University of Padua (protocol no. 4751) and were conducted according to the principles expressed in the Declaration of Helsinki.

### Experimental procedure

Children and their families were welcomed to the University’s hdEEG lab. Children were given time to familiarize themselves with the environment and with the experimenters while all experimental procedures were explained to parents, who provided written consent for their children’s participation. After assessing children’s oral assent to participate in the activities, a hdEEG 128-channel sensor net was applied.

The protocol included three main activities in the following order: 1) resting state session 1 (7 minutes), 2) resting state session 2 (7 minutes), 3) neuropsychological tests (∼10 minutes). Between each activity, children watched ∼ five minutes of an entertaining cartoon (The House of Mickey Mouse, Walt Disney). In the meantime, parents filled in a series of questionnaires assessing children’s behavioral and emotional profiles. Parental questionnaires were administered through the online platform *Qualtrics Surve*y (Qualtrics, Provo, UT).

### Cognitive measures

Non-verbal reasoning was assessed using the Coloured Progressive Matrices, in which children are required to complete a series of geometrical patterns and shapes (CPM, series A, B, AB; Raven & Court, 1938). Moreover, in order to obtain a wide descriptive profile of children’s CC abilities, we collected both neuropsychological measures and parental questionnaires assessing everyday behavior, executive functioning, and emotion regulation.

#### Neuropsychological measures

Phonological fluency was assessed as a measure of lexical access based on phonological cues and required children to say all the words they knew starting with a specific letter in one minute (BVN 5-11, Bisiacchi et al., 2005). Backward digit span (BDS) was assessed as a measure of verbal working memory and required children to retrieve a series of numbers in reverse order (BVN 5-11, Bisiacchi et al., 2005). Finally, gift wrap and gift wait (FE-PS 2-6, Usai et al., 2017) were administered to assess children’s levels of self-regulation and inhibition. Here, children were asked to wait with their eyes closed while the experimenter was wrapping a gift for them (total duration one minute); afterwards, the wrapped gift was placed in front of them and they were told that they could open it, but the longer they waited before opening (maximum waiting time four minutes), the bigger the gift would be.

**Table 1.** Cognitive tests. Here we report *means*, *standard deviations* and *range values* calculated for the neuropsychological tests across our sample. To derive a measure of phonological fluency, we calculated the mean number of words per category (total n° of words/ n° of categories) per participant; for Gift wrap and Gift time, we calculated the total time in seconds per participant. The sample’s scores were within the Italian normative range in all the reported measures.

| TEST | SCORE<br>*mean ± sd (range)* |
| --- | --- |
| Coloured Progressive Matrices | 20.0 ± 5.0 (10-34) |
| Phonological fluency | 2.3 ± 1.5 (0.7-7.3) |
| Backward digit span | 2.1 ± 1.2 (0-4) |
| Gift wrap (time in sec) | 46.9 ± 18.0 (1-60) |
| Gift wait (time in sec) | 98.6 ± 99.2 (0-240) |

#### Behavioral questionnaires

The behavioral and emotional profile of children was assessed using the Conners revised scales for parents (CPRS; Conners et al., 1998; Italian adaptation by Nobile et al., 2007), a parental questionnaire that comprises a series of statements about children’s behavior in everyday life. Children’s executive functioning in ecological contexts (e.g., home) was assessed using the Behavior Rating Inventory of Executive Function Preschool version (BRIEF-P; Gioia et al., 1996; Italian adaptation by Marano et al., 2014), a parental questionnaire that comprises a series of statements about children’s executive functioning in everyday life. Children’s emotion regulation was assessed using the Emotion Regulation CheckList (ERC; Molina et al., 2014), a parental questionnaire that comprises a series of statements assessing emotionality and regulation in children (e.g., affective lability, intensity). Finally, the Intolerance of Uncertainty scale for children (IUS-C, Comer et al., 2009; Italian adaptation by Bottesi et al., in prep.) was administered as a measure of cognitive adaptability in unpredictable contexts. Indeed, this parental questionnaire assesses the dispositional inability to tolerate the aversive reactions triggered by a perceived lack of sufficient and/or salient information.

**Table 2.** Parental questionnaire scores. Here we report *means*, *standard deviations* and *range values* calculated for the parental questionnaires across our sample. For each questionnaire, only total scales (highlighted in bold) entered subsequent statistical analyses. The sample’s scores were within the Italian normative range in all the reported questionnaires’ subscales.

| TEST | SCORE<br>*mean ± sd (range)* |
| --- | --- |
| Conners (Oppositivity) | 2.1 ± 1.3 (0-5) |
| Conners (Cognitive problems and disattention) | 4.5 ± 2.3 (0-9) |
| Conners (Hyperactivity) | 3.4 ± 2.2 (0-10) |
| Conners (Anxiety and shyness) | 2.4 ± 1.5 (0-6) |
| Conners (Perfectionism) | 2.4 ± 2.1 (0-8) |
| Conners (Social problems) | 2.1 ± 1.5 (0-6) |
| Conners (Psychosomatic problems) | 0.2 ± 0.7 (0-4) |
| Conners (ADHD index) | 3.4 ± 3.1 (0-12) |
| **Conners (DSM-IV total scale)** | **9.6 ± 5.3 (0-19)** |
| BRIEF (Inhibition) | 21.8 ± 3.3 (16-28) |
| BRIEF (Shifting) | 12.5 ± 2.6 (10-21) |
| BRIEF (Emotion regulation) | 13.9 ± 2.3 (10-18) |
| BRIEF (Working memory) | 21.0 ± 3.8 (17-31) |
| BRIEF (Planification-organization) | 13.7 ± 2.1 (11-19) |
| **BRIEF (Global Executive Composite)** | **82.8 ± 9.5 (69-111)** |
| ERC (Emotional regulation) | 27.0 ± 5.6 (0-32) |
| ERC (Lability/negativity) | 53.7 ± 9.7 (0-62) |
| **ERC (total scale)** | **80.7 ± 14.8 (0-94)** |
| **IUS-C (total scale)** | **23.3 ± 8.1 (0-42)** |

### hdEEG recording

The high spatial resolution EEG signal was recorded through a 128-channel Geodesic high-density EEG System (EGI® GES-300), with electrical reference to the vertex. A sampling rate of 500 Hz was used and impedance was kept below 60 kΩ for each electrode. Preprocessing was performed through EEGLAB v2022.0 (Delorme & Makeig, 2004).

## Data analysis

### hdEEG source reconstruction

#### Preprocessing and co-registration

The continuous EEG signal was first downsampled at 250 Hz and then bandpass-filtered (1-30 Hz) using a Hamming windowed sync finite impulse response filter. After filtering, the continuous EEG was visually inspected and bad segments (e.g., gross motor artifacts) were manually removed. Independent component analysis (ICA; Stone, 2002) using the Infomax algorithm (Bell & Sejnowski, 1995) was used to perform data cleaning. Independent components were visually inspected in topography and time-series, and those clearly related to eye blinks, eye movements, muscle artifacts and heartbeat were discarded. The remaining components were then projected back to the electrode space to obtain a cleaner EEG signal. Finally, flat and bad channels were reconstructed with the spherical spline interpolation method (Perrin et al., 1989). The data were then re-referenced to the average of all electrodes. Participants’ hdEEG data were co-registered using a natural (asymmetric) NIHPD Objective 1 scan template intended for preschool-aged children (4.5-8.5 years; Fonov et al., 2011). Co-registration was performed using the digitized scalp locations and fiducial markers using an iterative closest point algorithm in SPM12 (Ashburner et al., 2014). A forward model was fitted using the Boundary Element Method (BEM; Hall & Hall, 1994).

#### Source-localisation and parcellation

Final preprocessing steps were implemented using the OHBA Software Library (OSL v2.0.3; OHBA Analysis Group, 2017) and OHBA’s Hidden Markov Model Library (HMM-MAR; Vidaurre et al., 2016). First, a covariance matrix was computed across the whole-time course for each participant and PCA rank reduction allowed regularization of the obtained matrix to 50 dimensions. Then, a linearly-constrained minimum variance beamformer was used to estimate whole-brain source-space activity for points in an 8mm grid (Van Veen et al., 1997). Data dimensionality was reduced so that each individual brain activity was estimated as a series of time-courses for 3,559 source locations across the brain using signal-space separation algorithm (Woolrich et al., 2011). Afterwards, hdEEG data were further reduced into a 38-node cortical parcellation following the method proposed in Quinn et al. (2018). Finally, the parcellation was 13inarized to estimate a single time-course per node from the first principal components across voxels; crucially, this resulted in the reduction of individual time-courses to 38 parcels instead of 3,559 voxels, enabling additional corrections for signal leakage.

#### Additional preprocessing steps

Following OHBA’s HMM-MAR library, additional preprocessing steps were taken prior to the Hidden Markov Model initialisation: detrending, signal standardization and corrections for signal leakage. First, detrending removed linear trends in the data for each channel separately; second, participants’ concatenated time-courses were standardized. Next, signal leakage introduced by source reconstruction with zero temporal lag was corrected using multivariate orthogonalization (Colclough et al., 2015). Then, the Hilbert transform allowed absolute signal amplitude estimation for each source at each timepoint. See Figure 2 for a visual schematic representation of the preprocessing procedure.

**Figure 2.**
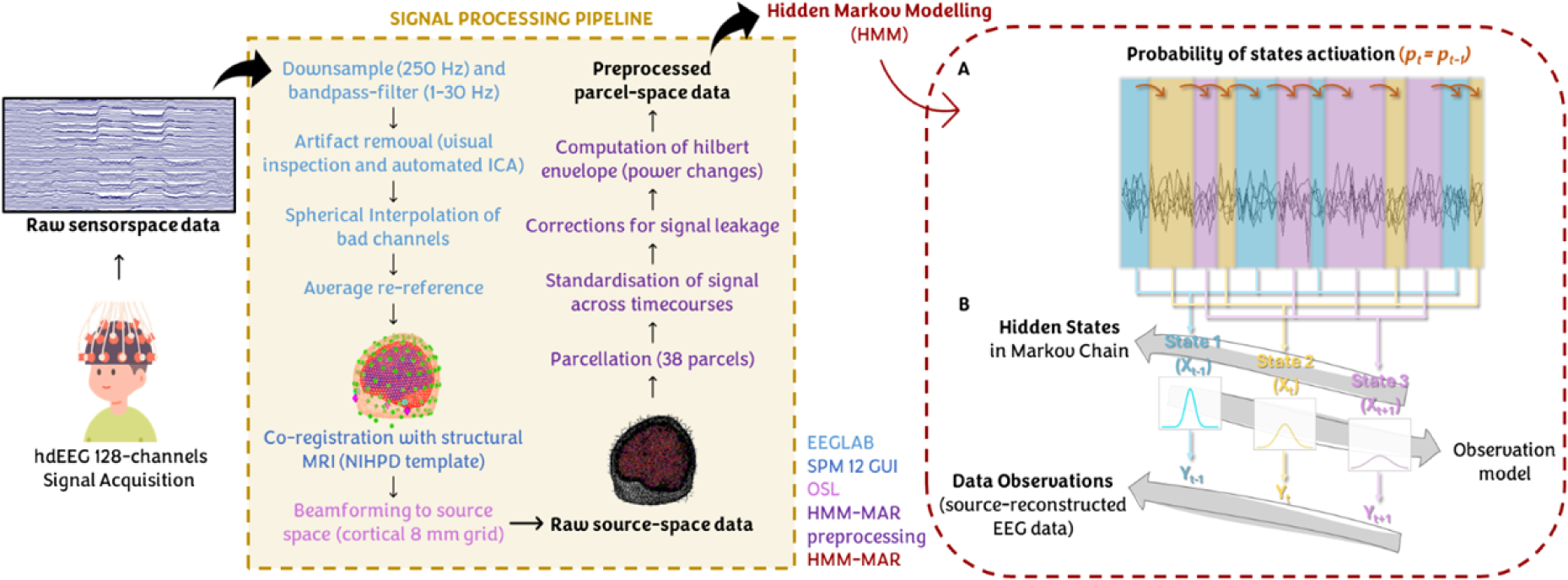
Data preprocessing pipeline and graphical representation of the HMM. On the left is a visual representation of each step of our hdEEG data preprocessing pipeline, in addition to the software packages and toolboxes used to complete each step of the pipeline. On the right is presented the HMM: A) the basic principle assumes that a time-series can be described using a hidden sequence of a finite number of states. Here, the time-series is partitioned into three states denoted by the blue, yellow and pink slabs. The model assumes that the probability (*p*) of each state at time point *t* depends on which state was active at time point *t-1*, as represented by the brown arrows. B) The model then assumes that the data observed in each state are drawn from a probabilistic observation model.

#### Hidden Markov Modeling (HMM)

In the present study, we used the HMM-MAR toolbox (Vidaurre et al., 2016), developed by the Oxford Centre for Human Brain Activity (OHBA) to infer a Hidden Markov Model from resting-state hdEEG time-series data. Hidden Markov Models (HMMs) refer to a set of unsupervised machine learning techniques that allow the segmentation of observed time-series data into a set of discrete hidden functional states (HMM states). These HMM states recur over millisecond timescales and are mutually-exclusive in time. A single model infers HMM states assuming they all have the same probabilistic distributions but with different parameterisations (Figure 2). In the present study, we employed the same formal definitions as in Zdorovtsova et al. (2023; see S1 in Supplementary Materials). The HMM-MAR toolbox computes a range of outputs that are useful in estimating different HMM states’ features. First, each state’s temporal characteristics were quantified in terms of state fractional occupancies (i.e., the fraction of the total time spent in a state), state lifetimes (i.e., the average time spent in a given state before transitioning to another state) and interval lengths (i.e., the average time it takes to re-enter a given state). Second, a switching rate (i.e., the frequency of state switching across an individual time-course) was calculated for each participant, providing a measure of individual network stability. The joint probabilities of transitions between pairs of states contained in the HMM output also allowed to compute state transition probability matrices for each participant, as well as for the entire concatenated time-course. Moreover, these allowed us to calculate an entropy rate estimate (i.e., a measure of average uncertainty generated by a transition within a sequence) per participant following the method employed by Zdorovtsova et al. (2023).

#### Analytic Plan

Here we provide a brief description of our research questions and the analyses we conducted to address them. Statistical analysis was performed using R (R Core Team, 2021; version 4.3.2). A Bayesian framework was used when fitting linear, generalized or multivariate linear models (R package: ‘brms’; Bürkner, 2017). Models’ specifications (i.e., number of chains, samples, burn-in, family distribution, custom or default priors) are specified in the Supplementary Materials. Convergence was assessed by examining the R-hat values (with a maximum accepted value for satisfactory convergence of 1.05 as suggested by Vehtari et al. 2021a), and by visual inspection of traces and the posterior predictive check. We employed leave-one out (LOO) cross-validation to evaluate the model performance and identify potential influential observations using k-Pareto diagnostics (R package: ‘loo’; Vehtari et al., 2021b). Based on the k-Pareto diagnostic plot, we considered observations with a Pareto shape parameter exceeding 0.7 as potentially influential. For each model, we report the estimated parameter value β, with an 89% Highest Density Interval (HDI) [lower limit, upper limit]. The use of an 89% HDI provides a narrower interval compared to traditional 95% intervals, thus allowing for a more focused estimate while still maintaining a high level of confidence. This choice of interval reflects a balance between precision and robustness in our inference (Makowski et al., 2019). Moreover, we report standardized betas (βs), computed as the ratio of the unstandardised beta coefficient to the standard deviation of the predictor variable (sigma), in order to offer a consistent measure of the relative impact of predictors on the outcome.

For hypothesis testing, we used the Region of Practical Equivalence (ROPE; Kruschke, 2018; R package: ‘bayestestR’, Makowski et al., 2019). The ROPE is defined as a region corresponding to the null value and the bulk of the posterior of a given parameter is compared with this region of values. If the HDI is completely outside the ROPE, the null hypothesis is ‘rejected’, while if the HDI overlaps with the ROPE the null hypothesis is ‘accepted’. For descriptive purposes, we consider less than 5% strong evidence of a relevant effect. We would like to point out that our observed variables are often very small, on a scale of thousandths; consequently, ROPE ranges tend to be generally narrow.

### Individual neural variability

First, two linear models were fitted separately for switching rate (SR) and entropy rates (see Table 3); specifically, each model comprised values of SR (standardized) or entropy (standardized) during the second resting state session as dependent variables and the values of SR or entropy during the first resting state session as independent variables. The models included a total of 39 observations (see Supplementary S2 for models’ specifications). Here, we used a ROPE range = [-0.2, 0.2] for equivalence testing.

**Table 3.**
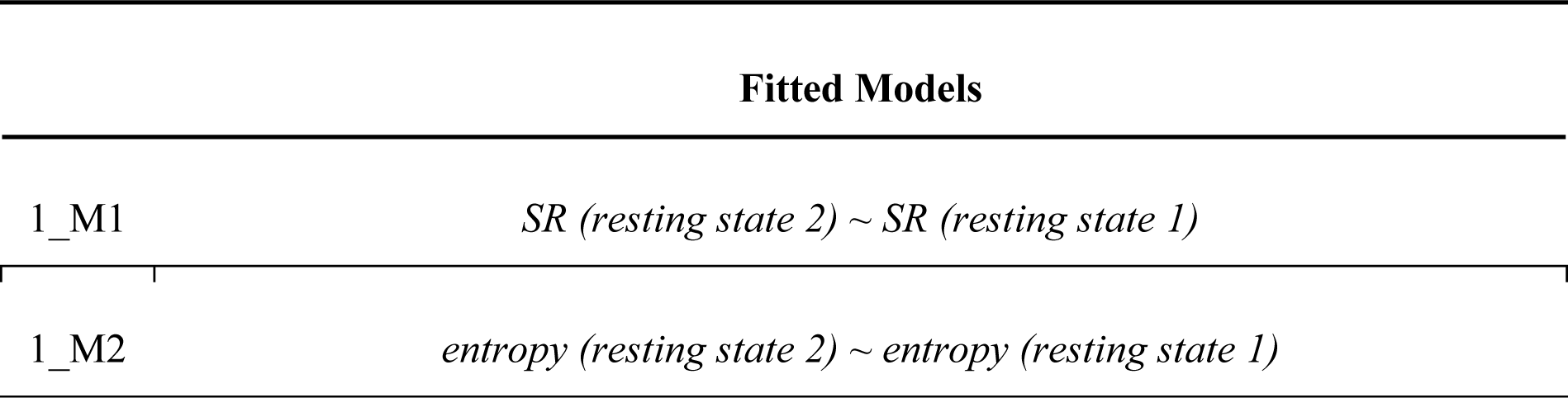
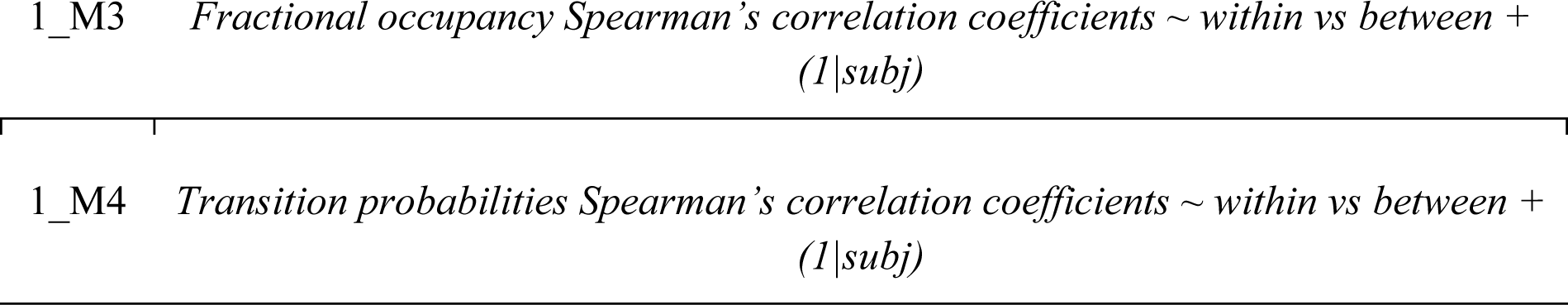
Fitted models to assess test-retest reliability of the HMM-derived indices.

Second, for fractional occupancies (N° states x N°subjects) and state transition probabilities (N°states^2 x N°subjects) we computed both within- and between-subjects Spearman correlations across the two resting state sessions. For both fractional occupancies and state transition probabilities, within-subjects correlations were computed as the correlation of each subjects’ measures between resting state 1 and resting state 2 (i.e., for each subject, 6 x 6 fractional occupancies and 36 x 36 transition probabilities; output = N°subjects x 1 correlation coefficient). Between-subjects correlations were computed as the correlation of each subject’s measures with the measures of each other subject at resting state 2 (i.e., for each subject, 6 x 6 x (N°subjects-1) fractional occupancies and 36 x 36 x (N°subjects-1) transition probabilities; output = N°subjects x (N°subjects-1) correlation coefficients). Afterwards, two separated generalized linear models were fitted for fractional occupancy and transition probabilities (see Table 3) with Spearman’s correlation coefficients as dependent variables, the type of correlation (within vs between) as independent variables and with a random intercept per subject. The model included a total of N°subjects x within correlation (one per subject) + N°subjects x between correlation (38 per subject) (see Supplementary S2 for model specifications). Here, we used a ROPE range = [-0.01, 0.01] for equivalence testing.

### Age- and gender-related differences

In the following analysis, we evaluated the effect of age (standardized, in months) and gender on the HMM indices computed for the first set of resting state recordings: 1) SR, 2) entropy rates, 3) states’ fractional occupancies and 4) probabilities of transitioning into each state (i.e., computed as the mean of transition probabilities toward each state, excluding self-transitions). This latter measure reduced data dimensionality (from 36 to 6 transitions) and allowed us to formulate more precise hypotheses. We fitted two separate linear models for SR and entropy rates and one multivariate linear model for transition probabilities. For fractional occupancies we run separate linear models for each state (see Table 4). The models included a total of 39 observations. Here we used a default ROPE range for equivalence testing (see Supplementary S3 for models’ specification).

**Table 4.** Fitted models to assess age and gender related effects on HMM-derived indices.

| Fitted Models |  |
| --- | --- |
| 2_M1 | SR ~ age (months) + gender |
| 2_M2 | entropy ~ age (months) + gender |
| 2_M3 | (Pr transition to State1 + Pr transition to State2 + ... + Pr transition to State6) ~ age (months) + gender |
| 2_M4 | FO State1 ~ age (months) + gender |
| 2_M5 | FO State2 ~ age (months) + gender |
| 2_M6 | FO State3 ~ age (months) + gender |
| 2_M7 | FO State4 ~ age (months) + gender |
| 2_M8 | FO State5 ~ age (months) + gender |
| 2_M9 | FO State6 ~ age (months) + gender |

### Individual neural variability and cognitive control

In the following analysis, for each subject we used HMM indices computed for the first set of resting state recordings (see above). To assess the relationship between cognitive and behavioral measures with HMM-derived indices, we fitted two multivariate linear models for each HMM index (each index was standardized) entering as dependent variables either 1) neuropsychological (NPS) measures (i.e., phonological fluency mean score, Gift wrap time and CPM score) or 2) questionnaires total scales (i.e., CPRS DSM-IV, BRIEF Global Executive Composite, ERC and IUS). Backward digit span and Gift wait were excluded from NPS models due to their non-normal distribution in order to maintain the assumption of normality necessary for reliable estimation. Age (standardized, in months) and gender entered as covariates in all the models (see Table 5). NPS models included a total of 39 observations, questionnaires’ models included a total of 38 observations due to missing data for one participant. Here we used a default ROPE range for equivalence testing (see Supplementary S4 for models’ specification).

**Table 5.** Fitted models to assess the relationship between cognitive control measures and HMM-derived indices.

| Fitted Models |  |
| --- | --- |
| 3_M1 | $(fluency, Wrap\ time, CPM) \sim SR + age\ (months) + gender$ |
| 3_M2 | $(fluency, Wrap\ time, CPM) \sim entropy + age\ (months) + gender$ |
| 3_M3 | $(fluency, Wrap\ time, CPM) \sim FO\ state1 + FO\ state2 + \dots + FO\ state6 + age\ (months) + gender$ |
| 3_M4 | $(fluency, Wrap\ time, CPM) \sim Pr\ transition\ to\ State1 + Pr\ transition\ to\ State2 + \dots + Pr\ transition\ to\ State6 + age\ (months) + gender$ |
| 3_M5 | $(CPRS, GEC, ERC, IUS) \sim SR + age\ (months) + gender$ |
| 3_M6 | $(CPRS, GEC, ERC, IUS) \sim entropy + age\ (months) + gender$ |
| 3_M7 | $(CPRS, GEC, ERC, IUS) \sim FO\ state1 + FO\ state2 + \dots + FO\ state6 + age\ (months) + gender$ |
| 3_M8 | $(CPRS, GEC, ERC, IUS) \sim Pr\ transition\ to\ State1 + Pr\ transition\ to\ State2 + \dots + Pr\ transition\ to\ State6 + age\ (months) + gender$ |

## Results

### State characteristics for the six-state HMM

A six-state HMM solution was selected as it revealed distinct spatiotemporal activity patterns while mitigating the redundancy associated with higher-state solutions (Figure 3). Each state-map displays the average activation profile of each parcel for the concatenated hdEEG dataset (for each subject, resting state 1 and resting state 2). State-specific activations are plotted as yellow/red. State 1 involves activations over prefrontal and temporo-parietal areas, resembling a prominent anterior DMN-like state. State 2 involves a lateralized pattern of activation in temporo-parietal regions of the cortex including the precuneus, and can be configured as a posterior DMN-like state. State 3 and 4 involve posterior regions including somatomotor areas and, respectively, occipito-temporal and occipito-parietal areas. Finally, state 5 and 6 display, respectively, prominent prefrontal and fronto-temporal activations.

**Figure 3.**
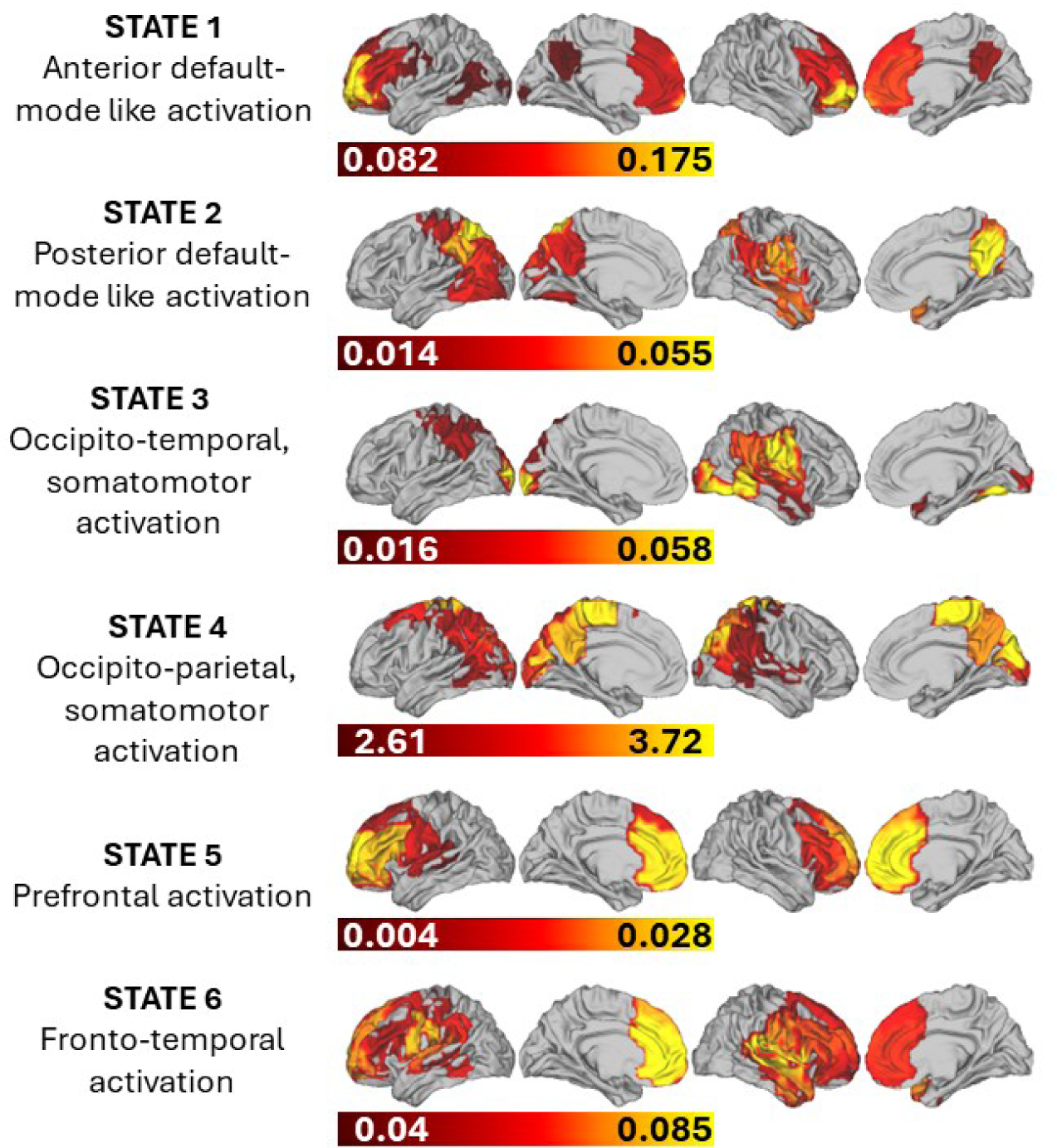
Six-state HMM. Here we present the visual representations of the six-state HMM. For each state, the top 60% of positive activations were plotted on a cortical surface using the HCP Workbench GUI. State labels correspond to our descriptions of the macroscopic features of the cortical activation pattern.

Temporal features of each HMM state were derived from state time-courses (Figure 4). Life times (LT; i.e., average temporal duration of state activation) and interval times (IT; i.e., average temporal duration between successive activations) show considerable variability within and between states, especially in states 4 and 5. In terms of fractional occupancies (FO; i.e., the total average time spent in each state), state 4 displays the lowest FO and state 5 the greatest variability. Means and standard deviations of LTs, ITs and FOs are summarised in Table 7.

**Figure 4.**
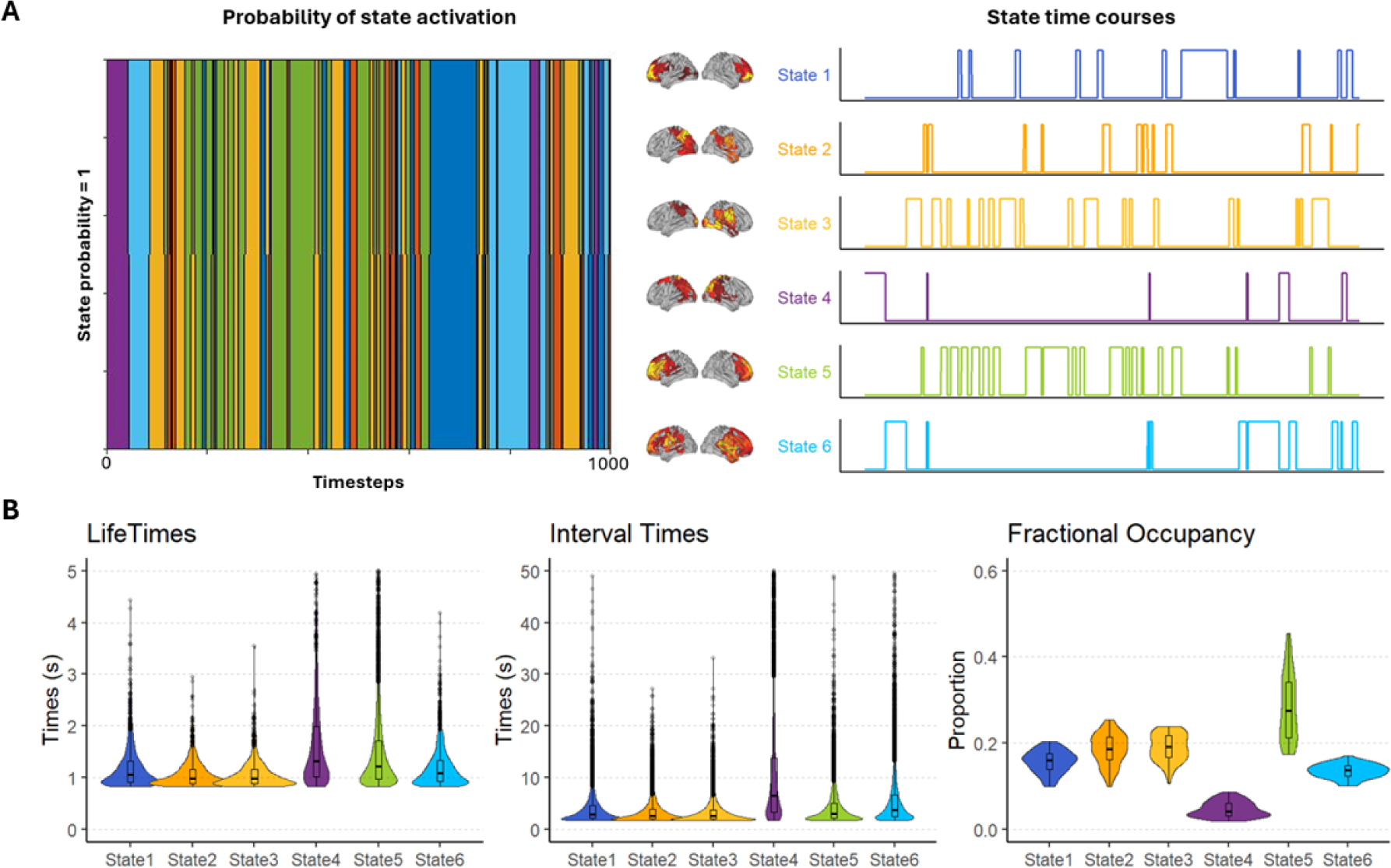
Temporal characteristics of the HMM states during both resting state sessions. A. The first 1000 timesteps (4 seconds sampled at 250Hz) of the Viterbi path (i.e., maximum a posteriori sequence of states in a HMM) and state time courses are presented. B. Life times (LT; left), interval times (IT; middle) and fractional occupancy (FO; right) of the states in our k = 6 HMM. Note that state LT and IT were thresholded respectively at 5s and 50s for visual representations.

**Table 7.** Means and standard deviations for state life times, interval times and fractional occupancy during both resting state sessions. Note that no thresholding was applied in the calculation of these metrics.

| **State** | **Life times (s)**<br>mean ± sd | **Interval times (s)**<br>mean ± sd | **Fractional**<br>**occupancy**<br>**(proportion)**<br>mean ± sd |
| --- | --- | --- | --- |
| State 1 (anterior DMN like) | 1.2 ± 0.4 | 4.0 ± 3.3 | 0.157 ± 0.026 |
| State 2<br>(posterior DMN like) | 1.1 ± 0.3 | 3.4 ± 2.3 | 0.187 ± 0.037 |
| State 3 (OT, SM) | 1.1 ± 0.3 | 3.4 ± 2.2 | 0.190 ± 0.032 |
| State 4 (OP, SM) | 1.7 ± 1.1 | 13.0 ± 16.4 | 0.047 ± 0.018 |
| State 5 (PF) | 1.6 ± 1.1 | 4.4 ± 3.8 | 0.284 ± 0.077 |
| State 6 (FT) | 1.2 ± 0.4 | 5.8 ± 5.9 | 0.136 ± 0.016 |

Finally, state transition probabilities between and across participants were computed. The dominant type of timepoint-to-timepoint transition was “self-transition”, forming the diagonal of the state transition probability matrix. Therefore, the diagonal was zeroed out to visualize state-to-state transition probabilities (Figure 5).

**Figure 5.**
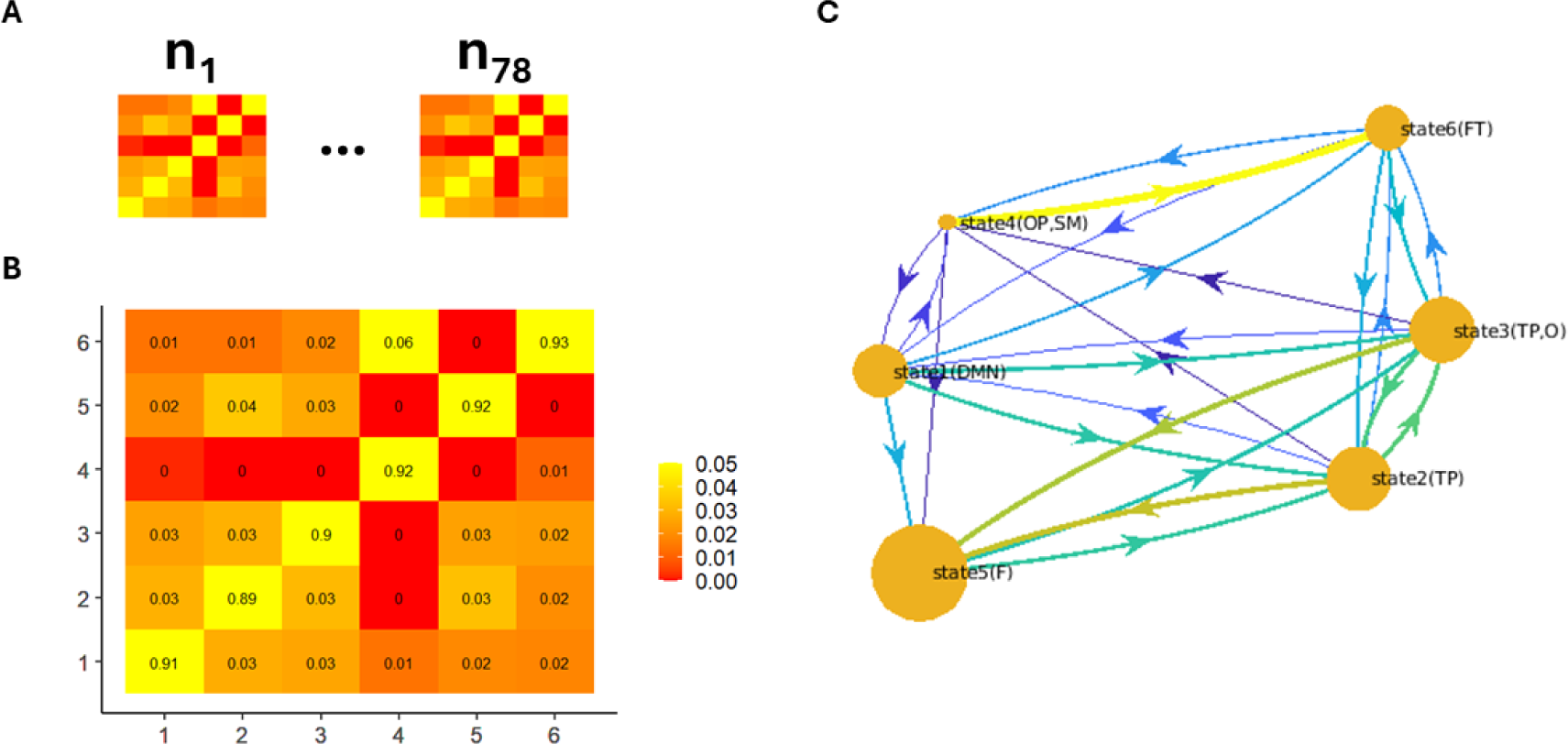
A. Joint posterior probabilities of state transitions for each subject were computed. B and C. Average transition probabilities across the entire sample during both resting state sessions (n = 78). Self-transitions were intentionally excluded for the purpose of plotting state-to-state transitions. No thresholds were applied. The node sizes in C reflect each state’s fractional occupancy.

### Individual switching rate, entropy, fractional occupancies and brain state transitions are reliable hallmarks of individual neural variability

To assess whether SR, entropy rates, fractional occupancies and brain state transitions are reliable hallmarks of individual neural variability, we ran separate linear and generalized linear models (see Table 3). For SR (see Table 3, 1_M1) the estimated regression coefficient (β) was 0.82 (89% HDI:[0.68, 0.98], βs = 1.46) and the percentage of β inside the ROPE was smaller than 5% (i.e., <1%). Similarly, for entropy (see Table 3, 1_M2), the estimated regression coefficient (β) was 0.82 (89% HDI:[0.66, 0.97], βs = 1.49) and the percentage of β inside the ROPE was less than 5% (i.e., <1%). Therefore, we concluded that both SR and entropy rates correlated between the two resting state sessions, suggesting that the dynamics of these states are relatively stable measures of individual differences.

For fractional occupancies (see Table 3, 1_M3), the estimated regression coefficient (β) was 0.08 (89% HDI:[0.04, 0.12], βs = 0.53) and the percentage of β inside the ROPE was less than 5% (i.e., <1%). Regarding transition probabilities (see Table 3, 1_M4), before computing within- and between-subjects Spearman correlations, for computational reasons we removed low-probability transitions (< 1 x 10^-5^) (N = 4, S2→S4, S2→S9, S3→S1, S3→S6). Indeed, non-zero variability is required to calculate correlations accurately. The estimated regression coefficient (β) was 0.03 (89% HDI:[0.02, 0.04], βs = 0.75) and the percentage of β inside the ROPE was less than 5% (i.e., <1%). Therefore, for both fractional occupancies and transition probabilities, results indicate greater correlation coefficients in the within-subjects compared to the between-subjects correlations (see Figure 6 for visual representation of the results). Overall, these results confirm within-subjects correlation of HMM-derived indices (i.e., SR, entropy rates, fractional occupancies and transition probabilities), supporting these as reliable hallmarks of individual neural variability.

**Figure 6.**
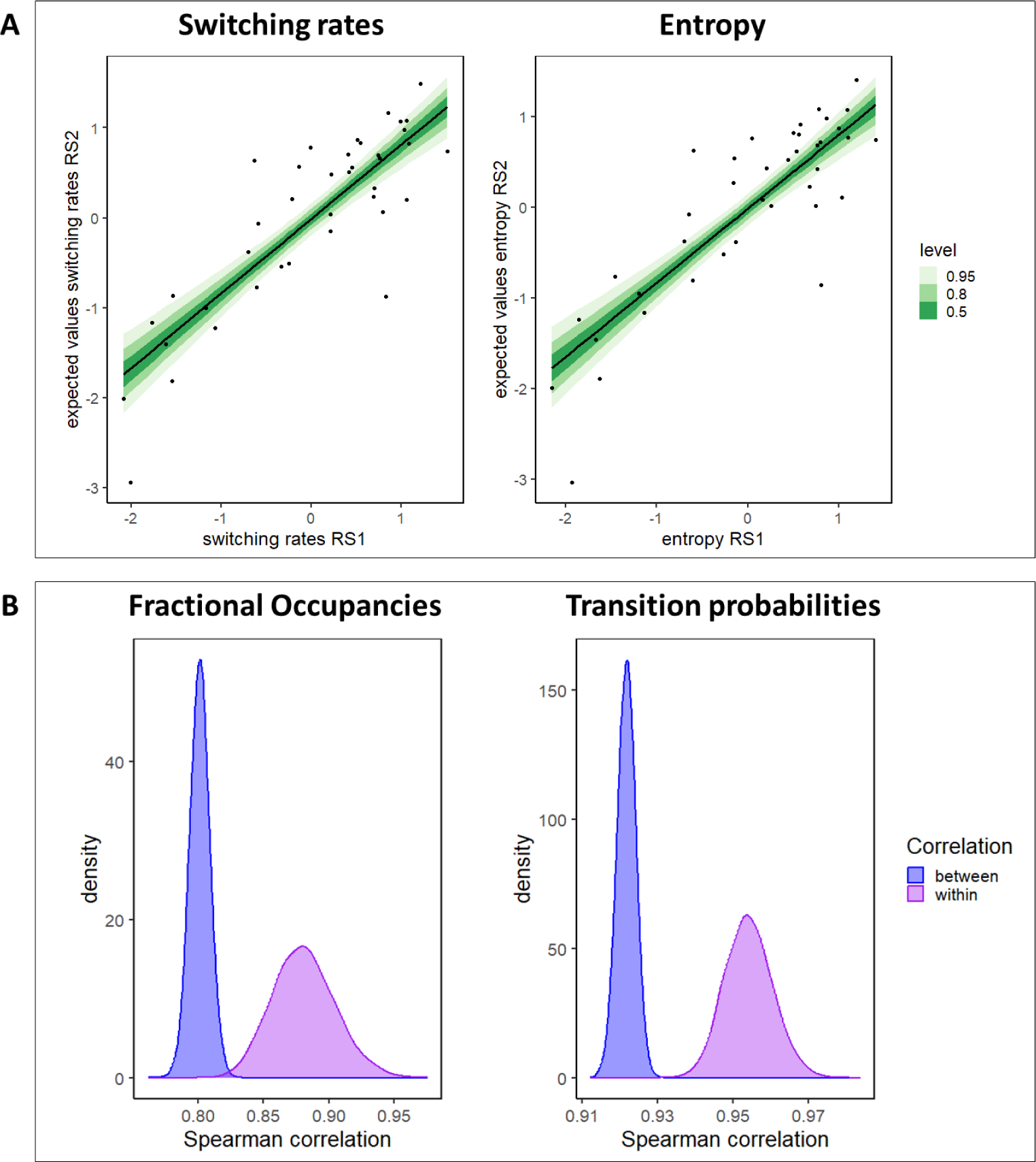
Panel A. The two figures display switching rates (left) and entropy (right) correlations between the two resting state sessions. On the X-axis are displayed expected values of switching rates/entropy in the first resting state session (RS1), on the Y-axis are displayed expected values of switching rates/entropy in the second resting state session (RS2). Panel B. The two figures display, for fractional occupancies (left) and transition probabilities (right), posterior density distributions (Y-axis) of Spearman’s correlation coefficients between the two resting state sessions (X-axis) for within (purple) and between (blue) correlations.

### Individual neural variability differs between boys and girls

To assess the effects of age and gender on HMM-derived individual neural variability indices we used separate linear and multivariate linear models (see Table 4). For both SR (see Table 4, 2_M1) and entropy rates (see Table 4, 2_M2), we found a main effect of gender, with boys displaying overall higher rates than girls (respectively, β = 0.01, 89% HDI:[0.00, 0.01], % inside ROPE < 1%, βs = 1 and β = 0.13, 89% HDI:[0.02, 0.23], % inside ROPE < 1%, βs = 0.65). Regarding transition probabilities (see Table 4, 2_M3), we found that boys have higher probability of switching into state 2 (β =0.002, 89% HDI:[0.000, 0.005], % inside ROPE < 3%, βs = 0.5) and a lower probability of switching into state 5 (β = −0.003, 89% HDI:[−0.005, - 0.000], % inside ROPE < 1%, βs = −0.6) compared to girls. Regarding fractional occupancies (see Table 4, 2_M4 – 2_M9), we found a main effect of gender in all the states, with boys spending overall more time in state 1 (β =0.02, 89% HDI:[0.00, 0.03], % inside ROPE < 1%, βs = 1), state 2 (β =0.02, 89% HDI:[0.01, 0.04], % inside ROPE < 1%, βs = 0.5), state 3 (β =0.03, 89% HDI:[0.01, 0.05], % inside ROPE < 1%, βs = 1) and state 6 (β =0.02, 89% HDI:[0.01, 0.02], % inside ROPE < 1%, βs = 2), and girls spending more time in state 4 (β =−0.02, 89% HDI:[−0.02, −0.01], % inside ROPE < 1%, βs = −1) and state 5 (β =−0.07, 89% HDI:[−0.11, −0.03], % inside ROPE < 1%, βs = −1). See Supplementary S3 for all the results, and Figure 7 for visual representation of the reported results. Overall, these results suggest the presence of gender-based but not age-based differences in individual neural variability, at least within this relatively narrow age band.

**Figure 7.**
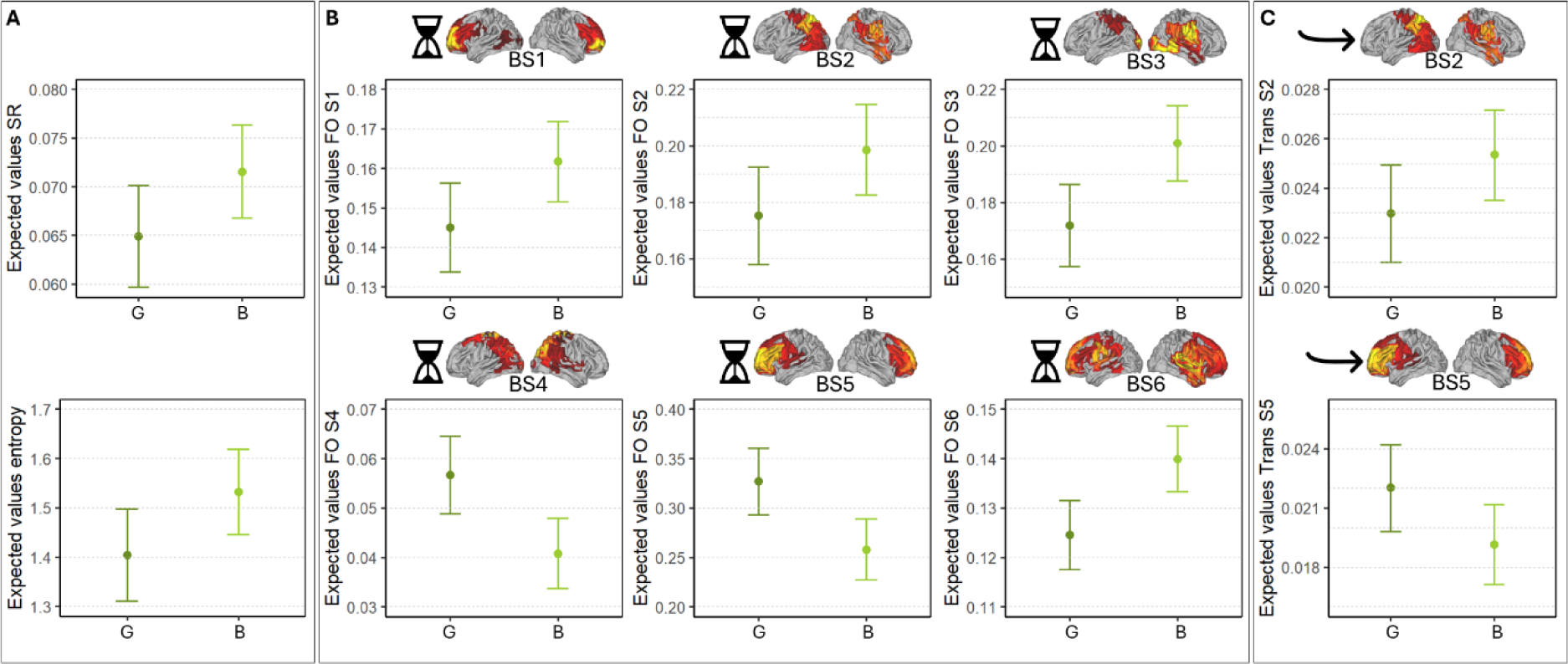
Gender-related differences in the HMM-derived indices. Panel A. The figures show on the y-axis expected values of switching (upper figure) and entropy (lower figure) rates, in boys (B; light green) and girls (G; dark green). Panel B. The figures show on the y-axis expected values of fractional occupancy (indicated symbolically by the hourglass) in brain state 1 (upper left), brain state 2 (upper center), brain state 3 (upper right), brain state 4 (bottom left), brain state 5 (bottom center) and brain state 6 (bottom right), in boys (B; light green) and girls (G; dark green). Panel C. The figures show on the y-axis expected values of transition probabilities (indicated symbolically by an arrow) of switching toward brain state 2 (upper figure) and state 5 (bottom figure), in boys (B; light green) and girls (G; dark green).

### Brain state transitions are related to preschoolers’ cognitive control

To unravel the relationship between individual neural variability and cognitive control we used separate multivariate models (see Table 5). For transition probabilities (see Table 5, 3_M8), we found that transitioning into brain state 2 is generally associated with worse behavioral and emotional profiles on the parental questionnaires: reduced Emotion Regulation CheckList (ERC) scores (β =-3.29, 89% HDI:[-5.96, −0.65], % inside ROPE < 1%, βs = −0.57), increased BRIEF-P Global Executive Composite (GEC) scores (β =5.78, 89% HDI:[1.52, 9.76], % inside ROPE < 1%, βs = 0.65) and increased Conners DSM-IV total scale (CPRS DSM-IV) scores (β =3.82, 89% HDI:[1.51, 6.12], % inside ROPE < 1%, βs = 0.79). Conversely, transitioning into brain state 3 was associated with better emotional regulation profiles as indicated by increased ERC scores (β =3.01, 89% HDI:[0.53, 5.55], % inside ROPE < 1%, βs = 0.53), and transitioning into brain state 6 was associated with better emotional regulation and behavioral profiles as indicated by increased ERC scores (β =2.83, 89% HDI:[0.98, 4.81], % inside ROPE < 1%, βs = 0.49), reduced GEC (β =-2.91, 89% HDI:[-5.73, −0.01], % inside ROPE < 10%, βs = −0.33) and CPRS DSM (β =-2.07, 89% HDI:[-3.68, −0.43], % inside ROPE < 2%, βs = −0.43) scores. Fractional occupancies (see Table 5, 3_M7), switching (see Table 5, 3_M5), and entropy rates (see Table 5, 3_M6) did not predict differences on any of the questionnaires’ scales (see Supplementary S5 for all the results, and Figure 8 for visual representation of the reported results). Finally, none of the individual neural variability indices predicted NPS measures (see Table 5, 3_M1 – 3_M4; see Supplementary S6 for all the results).

**Figure 8.**
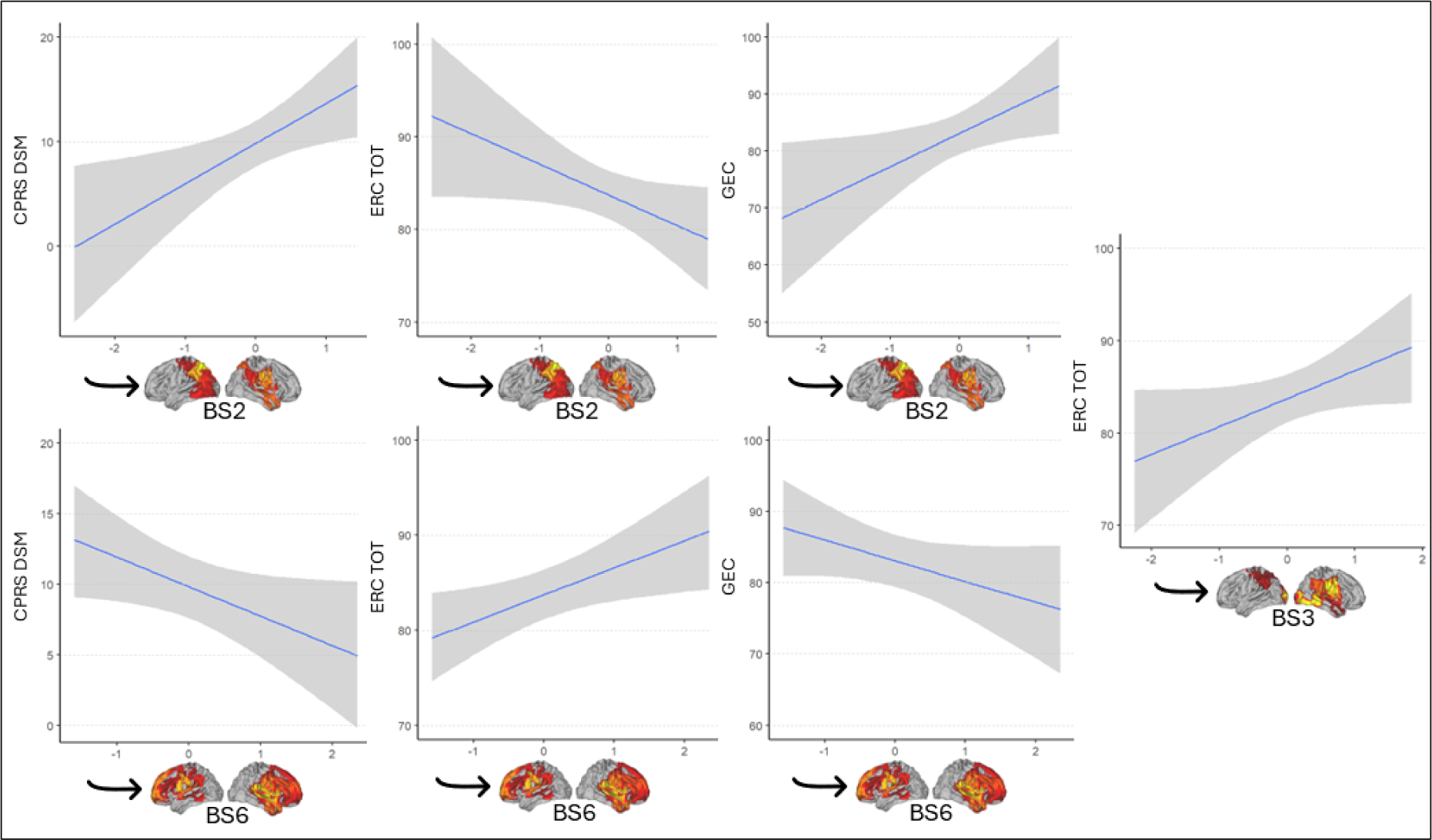
Transitions toward each state and questionnaire scores. The figures reported in the panel show on x-axis the expected values (standardized) of brain state transitions (indicated symbolically by an arrow) toward a certain brain state (i.e., BS2 = brain state 2, BS3 = brain state 3, BS6 = brain state 6) and on the y-axis the expected values of the questionnaires’ scales (GEC = global executive composite (BRIEF-P), ERC_tot = emotion regulation checkList total scale, CPRS_DSM_tot = conners DSM-IV total scale) scores.

Overall, results only partly confirm that transitioning toward CCN-like states predicts better behavioral and emotional regulation profiles as assessed using parental questionnaires.

## Discussion

In the present study, we investigated preschoolers’ resting-state neural dynamics by inferring a six-state Hidden Markov Model from hdEEG data. Our model identified discrete spatiotemporal patterns mimicking well known RSNs, including the anterior and posterior default-mode, temporo-parietal, occipital, sensorimotor, frontal and fronto-temporal networks. The states were characterized by short (< 150 ms) time courses, supporting the idea of RSNs as a system of rapid synchronization and switching dynamics. Individual switching rates, entropy rates, fractional occupancies and state transition probabilities were reliable across different recording sessions, supporting HMM inference as a robust methodology to investigate individual neural variability in the developing brain. This is particularly relevant considering the well-known difficulties in collecting good-quality neural data from young populations. Assessing the reliability of these neural indices is a fundamental step to study how they relate to individual differences in cognitive functioning. Our analyses served as an attempt to describe how different neural indices of brain dynamics are intrinsically related to broader measures of behavior and cognition in preschoolers. Overall, our results indicate that HMM-derived metrics show gender-related differences and predict CC measures. We discuss these results further in the following sections.

### Gender, but not age, explains differences in individual neural hallmarks

We observed gender-related differences in almost all the HMM-derived indices. This is in line with previous studies that found differences in static (Ritchie et al., 2018) and dynamic (Scofield et al., 2019) functional connectivity between girls and females. For instance, Scofield et al. (2019) found that school-aged boys show greater dwell time in states related to the ventral attention network, DMN and somatomotor network, before switching to another state. In a similar vein, our analysis revealed that, compared to girls, boys tend to switch more frequently between states, especially toward states that might be considered task-negative or default-mode like (state 2 involves activations over the precuneus, which is reminiscent of a posterior default-mode activation; Raichle, 2015) and spend on average more time in almost all the states including state 1 (anterior default-mode like). Conversely, girls switch less frequently between states but mostly toward CCN-like states, where they spend more time: state 5 (prefrontal) and state 4 (occipito-parietal, somatomotor). Interestingly, both prefrontal and parietal areas have been proposed as two core hubs underlying high-order cognitive functioning. For instance, individual differences in reasoning and working memory abilities might rely on the interplay between parietal and frontal association cortices, as well as other structural and functional parameters like their volume, gray matter density and mean diffusivity (Gur et al., 2021; Jung & Haier, 2007; Li & Tian, 2014). Additionally, the fronto-parietal network is considered a modal controllability hub, which is the ability of a given network to drive the whole brain into states that are difficult to reach, such as those entered under high cognitive demand (Gu et al., 2015). Greater engagement (fractional occupancy) of these states in girls might explain their reduced switching and entropy rates compared to boys. Therefore, we speculate that these gender-related differences in brain dynamics may heighten boys neurobiological risk of atypical developmental trajectories associated with compromised CC (Bölte et al., 2023). Contrary to our expectations (Kupis et al., 2021) we did not find age-related differences in the HMM-derived indices. This could be due to the restricted age-range considered in the present study (i.e., 4-6 years).

### Brain state transitions are associated with cognitive control

The preschool period is a sensitive window for the development of CC, particularly due to increasing patterns of connectivity between the prefrontal cortex, the anterior cingulate cortex, and the parietal cortex (Fiske & Hombloe, 2019). In the present study, we found that specific resting-state patterns of brain state transitions are associated with cognitive control, as assessed using parental questionnaires. More specifically, we found that a higher probability of transitioning into state 6 (fronto-temporal) was associated with better scores in questionnaires assessing executive functioning in everyday life (BRIEF-P; Gioia et al., 1996), behavioral difficulties (CPRS; Conners et al., 1998) and emotion regulation abilities (ERC; Molina et al., 2014). On the contrary, we found that a higher probability of transitioning into state 2 (posterior default-mode like) was associated with worse scores in the same measures. Overall, these findings build on previous studies supporting the important role of the prefrontal cortex for CC development (Friedman & Robbins, 2021). Moreover, it confirms and extends previous findings suggesting the relevance of brain dynamics for CC within the preschooler period (Cabral et al., 2017; Kupis et al., 2021). Specifically, previous evidence showed that adults performing differently in CC tasks displayed different brain state transitions (Cabral et al., 2017) and that, generally, brain state transitions relate to subject-specific cognitive traits (Vidaurre et al., 2017). Within this framework, our findings suggest that a higher tendency to enter prefrontal states might favor the engagement of CC abilities in response to external demands (Emili Balaguer-Ballester et al., 2011).

Interestingly, we also found that a higher probability of transitioning into state 3 (occipito-temporal, somatomotor), was associated with better emotional regulation (ERC; Molina et al., 2014). State 3 activation involves areas associated with the ventral circuit (fusiform area, occipital extrastriate and superior temporal sulcus) along with right frontal areas and might overlap with the social pathway recently proposed by Pitcher & Ungerleider (2021). This latter network is specialized for processing dynamic aspects of social perception. Therefore, we can hypothesize a relationship between the at-rest functional activation of this network and the way young children manage their emotion regulation in social contexts. Nevertheless, this hypothesis is speculative and would require further investigation.

Overall, it is interesting to note that we found relationships between state transition probabilities and parental questionnaires scores, but no relationship with direct cognitive measures. This might depend on the highly contextual-dependent performance of young children and, in turn, on the low test-retest reliability of such neuropsychological measures, which may not be sensitive enough to individual differences (Karalunas et al., 2020). Consequently, upcoming studies should aim to find more sensitive measures for assessing cognitive abilities in preschoolers.

Notably, in contrast to previous studies on adults (Nomi et al. 2017; Taghia et al., 2018; Vidaurre et al., 2017), and older school-aged children (Zdorovtsova et al., 2023), we only found state transition probabilities to be associated with CC, while we found no relationship with other HMM-derived neural indices (switching, entropy rates, and fractional occupancies). One potential explanation is that state transition patterns might represent a prominent and more sensitive feature of CC development during the preschool-age period compared to other neural indices. Indeed, during this time the brain undergoes substantial functional modifications, including changes in inter-regional activity (Brown & Jernigan, 2012), that may be better captured by state transitions rather than the other HMM indices. Future studies should address this point using different CC measures. Finally, task-evoked HMM-derived indices might provide a complementary view (Medaglia et al., 2018).

## Conclusions

We inferred a six-state Multivariate Gaussian Hidden Markov Model using resting-state hdEEG brain activity in a developmental, preschool-aged sample (4-6 years). These brain state characteristics derived from the HMM were reliable across different recording sessions, allowing their use as individual neural hallmarks, and differed between boys and girls. Moreover, transition patterns between states were predictive of individual differences on cognitive control measures (parental questionnaires scores). In line with previous evidence, the current study supports the importance of resting-state brain dynamics as an important scaffold for behavior and cognition. Moreover, for the first time, we extend these findings to the preschool period and suggest that brain state transitions might be particularly salient during this developmental window. Therefore, brain state transitions should be targeted by future studies investigating neurodevelopmental trajectories and early markers of neurodivergent development.

## Acknowledgments

We are grateful to all the participating children and their families. A special thanks goes to Martina Rizzuti, Kim Lai Cangiulli and Cristian Fanzolato for helping with data collection. We would also like to thank the Istituto Comprensivo “G. Santini” in Noventa Padovana (Padua, Italy) and “S.P.E.S.” (Padua, Italy) for the collaboration in participant recruitment. We also thank Prof. Joseph Onto for providing us with food for thought.

## Funding

This work was supported by Ricerca Corrente 2024 to Gian Marco Duma funds for biomedical research of The Italian Health Ministry. D.E.A. is supported by Medical Research Council Programme Grant MC-A0606-5, the Gnodde Goldman Sachs endowed Professorship in Neuroinformatics, and by the Templeton World Charity Foundation, Inc. (funder DOI 501100011730) under grant TWCF-2022-30510. D.E.A. and G.E. were supported by The James S. McDonnell Foundation Opportunity Award. All research at the Department of Psychiatry at the University of Cambridge is supported by the National Institute for Health and Care Research Cambridge Biomedical Research Centre (NIHR203312) and the NIHR Applied Research Collaboration East of England. Open access funding provided by Bibliosan.

## Conflict of Interest Statement

The authors declare no competing interests.

## Data Availability Statement

The data and analysis code that support the findings of this study will be made openly available in Open Science Framework (OSF) and can be accessed at: https://osf.io/agu27/?view_only=1fcfac943c89434e8f900968bb09b77d

## Notes

https://osf.io/agu27/?view_only=1fcfac943c89434e8f900968bb09b77d

### Competing Interest Statement

The authors have declared no competing interest.

