## Supplementary material for "Dynamic transient brain states in preschoolers mirror parental report of behavior and emotion regulation"

### Supplementary Materials

#### S1 - Hidden Markov Modeling (HMM)

In the present study we used the same formal definitions as in Zdorovtsova et al. (2023). Specifically, we assumed a temporal dependency of the states (i.e., the probability of each state activation at each time point depends on the state active in the previous time point) and a multivariate Gaussian observational model (i.e., the observation model links the observed time-series with the hidden HMM states assuming that each state has a specific mean and covariance). Participants' time-series data was concatenated before running the HMM in order to retrieve both group-level estimates of the states and individual time-courses of the states.

#### S2 - Are individual switching rate, entropy, fractional occupancies and transitions probabilities reliable hallmarks of individual neural variability?

All the following models were run using 4 chains, with a total of 8,000 iterations, discarding the initial 4,000 iterations as burn-in.

**Switching rate**

*Model switch:*

The model was run using normal (0, 1) prior Intercept and student t (3, 0, 5) for β parameter, with skew normal family distribution. Default priors were used for standard deviations (sigma: student t (3, 0, 2.5)) and alpha (normal (0, 4)).

*SR (resting state 2) ~ SR (resting state 1)*

**Entropy**

*Model entropy:*

The model was run using normal (0, 1) prior Intercept and student t (3, 0, 5) for β parameter, with skew normal family distribution. Default priors were used for standard deviations (sigma: student t (3, 0, 2.5)) and alpha (normal (0, 4)).

*Entropy (resting state 2) ~ Entropy (resting state 1)*

**Transition probabilities**

*Model transitions:*

The model was run using normal (0.9, 1) prior for Intercept with lower bound = 0 and upper bound = 1 and student t (3, 0, 5) for β parameter, with skew normal family distribution. Default priors were used for standard deviations (sigma: student t (3, 0, 2.5)), alpha (normal (0, 4)) and parameters standard deviations (sd: student t (3, 0, 2.5).

*spearman correlation coefficients ~ 1 + condition* (factor variable, 2 levels: within vs between) + *(1|subj)*

**Fraction occupancies**

*Model FO:*

The model was run using normal (0.9, 1) prior for Intercept with lower bound = 0 and upper bound = 1 and student t (3, 0, 5) for β parameter, with skew normal family distribution. Default priors were used for standard deviations (sigma: student t (3, 0, 2.5)), alpha (normal (0, 4)) and parameters standard deviations (sd: student t (3, 0, 2.5).

*spearman correlation coefficients ~ 1 + condition* (factor variable, 2 levels: within vs between) + *(1|subj)*

#### S3 - Are there age and gender related differences in individual neural variability?

**Switching rate**

The model was run using 4 chains, with a total of 10,000 iterations, discarding the initial 5,000 iterations as burn-in. For equivalence testing we used the default ROPE range computed by the rope_range function.

*SR ~ 1 + age (months) + gender*

The model was run using normal (0, 1) with lower bound for Intercept, student t (3, 0, 5) for β parameter and gaussian family distribution. Default priors were used normal (0, 4) for alpha and student t (3, 0, 2.5) for sigma.

| **Parameter** | **Coefficient** | **89% HDI** | **ROPE range** | **% ROPE** |
|---|---|---|---|---|
| Age (months) | -0.001 | [-0.004, 0.002] | -0.001, 0.001 | 42.53 % |
| Gender | 0.01 | [0.00, 0.01] | -0.001, 0.001 | 0.83 % |

**Entropy rate**

The model was run using 4 chains, with a total of 10,000 iterations, discarding the initial 5,000 iterations as burn-in. For equivalence testing we used the default ROPE range computed by the rope_range function.

$$entropy \sim 1 + age\ (months) + gender$$

The model was run using normal (0, 1) with lower bound for Intercept, student t (3, 0, 5) for β parameter and gaussian family distribution. Default priors were used normal (0, 4) for alpha and student t (3, 0, 2.5) for sigma.

| Parameter | Coefficient | 89% HDI | ROPE range | % ROPE |
|---|---|---|---|---|
| Age (months) | -0.02 | [-0.07, 0.03] | -0.02, 0.02 | 43.56 % |
| Gender | 0.13 | [0.02, 0.23] | -0.02, 0.02 | 0.00 % |

**Transition probabilities**

The model was run using 4 chains, with a total of 20,000 iterations, discarding the initial 10,000 iterations as burn-in. For equivalence testing we used the default ROPE range computed by the rope_range function.

$$(Pr\ transition\ to\ State\ 1,\ Pr\ transition\ to\ State\ 2\ \ldots\ Pr\ transition\ to\ State\ 6) \sim$$
$$1 + age\ (months) + gender$$

The model was run using normal (0, 1) with lower bound for Intercept and student t (3, 0, 5) for β parameter. Default priors were used student t (3, 0, 2.5) for sigma and lkj corr cholesky (1). Gaussian family distribution and the set_rescor parameter was enabled or set to TRUE to account for residual correlations.

| Response variable | Parameter | Coefficient | 89% HDI | ROPE range | % ROPE |
|---|---|---|---|---|---|
| **Pr transition to S1** | Age (months) | 0.0003 | [ -0.001, 0.001] | -0.0004, 0.0004 | 46.26 % |
| | Gender | 0.001 | [ -0.001, 0.003] | -0.0004, 0.0004 | 17.00 % |
| **Pr transition to S2** | Age (months) | -0.0003 | [-0.0014, 0.001] | -0.0004, 0.0004 | 47.92 % |
| | Gender | 0.002 | [0.0002, 0.005] | -0.0004, 0.0004 | 2.35 % |
| **Pr transition to S3** | Age (months) | -0.0001 | [-0.0012, 0.001] | -0.0004, 0.0004 | 49.81 % |
| | Gender | 0.001 | [-0.0011, 0.0032] | -0.0004, 0.0004 | 20.34 % |
| **Pr transition to S4** | Age (months) | -0.001 | [-0.0010, -0.0001] | -0.0002, 0.0002 | 6.64 % |
| | Gender | -0.001 | [-0.002, 0.0002] | -0.0002, 0.0002 | 14.35 % |
| **Pr transition to S5** | Age (months) | -0.0001 | [-0.0013, 0.0011] | -0.0004, 0.0004 | 50.72 % |
| | Gender | -0.003 | [-0.01, -0.0004] | -0.0004, 0.0004 | 0.00 % |
| **Pr transition to S6** | Age (months) | -0.001 | [-0.0014, 0.0002] | -0.0003, 0.0003 | 20.52 % |
| | Gender | 0.001 | [-0.0004, 0.003] | -0.0003, 0.0003 | 12.37 % |

**Fractional occupancies**

For fractional occupancies, we run separate models for each state instead of a single multivariate model because of convergence issues. The models were run using 4 chains, with a total of 10,000 iterations, discarding the initial 5,000 iterations as burn-in. For equivalence testing we used the default ROPE range computed by the rope_range function.

*FO State 1 ~ 1 + age (months) + gender*

*FO State 2 ~ 1 + age (months) + gender*

*FO State 3 ~ 1 + age (months) + gender*

*FO State 4 ~ 1 + age (months) + gender*

*FO State 5 ~ 1 + age (months) + gender*

*FO State 6 ~ 1 + age (months) + gender*

The models were run using normal (0, 1) with lower bound for Intercept, student t (3, 0, 5) for β parameter and gaussian family distribution. Default prior was used student t (3, 0, 2.5) for sigma.

| Response variable | Parameter | Coefficient | 89% HDI | ROPE range | % ROPE |
|---|---|---|---|---|---|
| **FO S1** | Age (months) | 0.01 | [-0.001, 0.01] | -0.002, 0.002 | 21.37 % |
| | Gender | 0.02 | [0.005, 0.03] | -0.002, 0.002 | 0.00 % |
| **FO S2** | Age (months) | 0.0004 | [-0.01, 0.01] | -0.004, 0.004 | 50.71 % |
| | Gender | 0.02 | [0.01, 0.04] | -0.004, 0.004 | 0.00 % |
| **FO S3** | Age (months) | -0.001 | [-0.01, 0.01] | -0.003, 0.003 | 52.87 % |
| | Gender | 0.03 | [0.01, 0.05] | -0.003, 0.003 | 0.00 % |
| **FO S4** | Age (months) | -0.002 | [-0.01, 0.002] | -0.002, 0.002 | 45.23 % |

| | | | | | |
|---|---|---|---|---|---|
| | Gender | -0.02 | [-0.02, -0.01] | -0.002, 0.002 | 0.00 % |
| **FO S5** | Age (months) | -0.01 | [-0.02, 0.01] | -0.008, 0.008 | 49.89 % |
| | Gender | -0.07 | [-0.11, -0.03] | -0.008, 0.008 | 0.00 % |
| **FO S6** | Age (months) | 0.003 | [-0.001, 0.01] | -0.002, 0.002 | 22.85 % |
| | Gender | 0.02 | [0.01, 0.02] | -0.002, 0.002 | 0.00 % |

#### S4 - Is individual neural variability functionally related to cognitive control?

All the following models were run using 4 chains, with a total of 10,000 iterations, discarding the initial 5,000 iterations as burn-in.

*Neuropsychological measures:*

All the models were run using normal (2, 15) with lower bound = 0 prior Intercept for phonological fluency, normal (48, 30) with lower bound = 0 for Wrap time Intercept and normal (20, 15) with lower bound = 0 for CPM intercept. For all the β parameters student t (3, 0, 3) was used. Default priors were used for correlated residuals (Lrescor: lkj corr cholesky(1)) and standard deviations (sigma: phonological fluency student t (3, 0, 2.5), CPM student t (3, 0, 4.4), Wrap time student t (3, 0, 3). Gaussian family distribution and the set_rescor parameter was enabled or set to TRUE to account for residual correlations.

*Questionnaires:*

All the models were run using normal (83, 17) with lower bound = 0 prior Intercept for ERC total, normal (83, 20) with lower bound = 0 for GEC Intercept, normal (10, 15) with lower bound = 0 for CPRS DSM intercept and normal (24, 17) for IUS Intercept. For all the β parameters student t (3, 0, 3) was used for all the β parameters, for fractional occupancies student t (3, 0 , 2) was used. Default priors were used for correlated residuals (Lrescor: lkj corr cholesky(1)) and standard deviations (sigma: CPRS DSM student t (3, 0, 5.2), ERC student t (3, 0, 8.2), GEC student t (3, 0, 8.9), IUS student t (3, 0, 8.9)). Gaussian family distribution and the set_rescor parameter was enabled or set to TRUE to account for residual correlations.

#### S5 - Individual neural variability and questionnaires

For equivalence testing we used the default ROPE range computed by the rope_range function. Utilizing the default ROPE calculation ensures consistency and objectivity in defining a practical range of parameter values, improving results comparability and aiding in the clear interpretation of effects within the analysis.

**Switching rates and questionnaires**

*(CPRS, GEC, ERC, IUS) ~ SR + age (months) + gender*

| Response variable | Parameter | Coefficient | 89% HDI | ROPE range | % ROPE |
|---|---|---|---|---|---|
| **CPRS** | SR | 1.08 | [-0.24, 2.48] | -0.54, 0.54 | 24.43 % |
| | Age (months) | -0.42 | [-1.69, 0.94] | -0.54, 0.54 | 46.34 % |
| | Gender | -0.15 | [-2.44, 2.16] | -0.54, 0.54 | 31.07 % |
| **GEC** | SR | 0.56 | [-1.80, 2.74] | -0.96, 0.96 | 49.55 % |
| | Age (months) | 0.08 | [-2.06, 2.34] | -0.96, 0.96 | 54.10 % |
| | Gender | -0.16 | [-3.60, 3.40] | -0.96, 0.96 | 37.19 % |
| **ERC** | SR | 1.24 | [-0.39, 2.99] | -0.66, 0.66 | 26.03 % |
| | Age (months) | -2.28 | [-2.23, 1.01] | -0.66, 0.66 | 43.39 % |
| | Gender | -0.62 | [-5.21, 0.69] | -0.66, 0.66 | 14.54 % |
| **IUS** | SR | -0.34 | [-2.29, 1.51] | -0.73, 0.73 | 47.48 % |
| | Age (months) | -0.71 | [-2.48, 1.23] | -0.73, 0.73 | 42.06 % |
| | Gender | 0.83 | [-2.29, 4.10] | -0.73, 0.73 | 28.83 % |

**Entropy rates and questionnaires**

*(CPRS, GEC, ERC, IUS) ~ entropy + age (months) + gender*

| Response variable | Parameter | Coefficient | 89% HDI | ROPE range | % ROPE |
|---|---|---|---|---|---|
| **CPRS** | Entropy | 1.12 | [-0.24, 2.50] | -0.54, 0.54 | 21.66 % |
| | Age (months) | -0.42 | [-2.64, 2.03] | -0.54, 0.54 | 49.48 % |
| | Gender | -0.19 | [-1.75, 0.87] | -0.54, 0.54 | 32.46 % |
| **GEC** | Entropy | 0.56 | [-1.76, 2.81] | -0.96, 0.96 | 53.16 % |
| | Age (months) | 0.07 | [-2.12, 2.33] | -0.96, 0.96 | 58.07 % |
| | Gender | -0.17 | [-3.60, 3.41] | -0.96, 0.96 | 38.52 % |
| **ERC** | Entropy | 1.16 | [-0.52, 2.85] | -0.66, 0.66 | 29.39 % |
| | Age (months) | -0.64 | [-2.29, 0.97] | -0.66, 0.66 | 46.19 % |
| | Gender | -2.30 | [-5.14, 0.82] | -0.66, 0.66 | 14.90 % |
| **IUS** | Entropy | -0.31 | [-2.23, 1.62] | -0.73, 0.73 | 49.78 % |
| | Age (months) | -0.70 | [-2.44, 4.05] | -0.73, 0.73 | 44.48 % |
| | Gender | 0.84 | [-2.65, 1.13] | -0.73, 0.73 | 30.15 % |

**Fractional occupancies and questionnaires**

*(CPRS, GEC, ERC, IUS) ~ FO state1 + FO state2 + … + FO state6 + age (months) + gender*

| Response variable | Parameter | Coefficient | 89% HDI | ROPE range | % ROPE |
|---|---|---|---|---|---|
| **CPRS** | FO State 1 | -0.27 | [-2.72, 2.26] | -0.54, 0.54 | 30.72 % |
| | FO State 2 | 1.82 | [-1.04, 4.67] | -0.54, 0.54 | 17.10 % |
| | FO State 3 | -1.63 | [-4.43, 1.26] | -0.54, 0.54 | 17.81 % |
| | FO State 4 | -1.76 | [-5.58, 2.40] | -0.54, 0.54 | 16.37 % |
| | FO State 5 | 0.51 | [-4.22, 5.06] | -0.54, 0.54 | 17.24 % |
| | FO State 6 | -0.50 | [-2.44, 1.42] | -0.54, 0.54 | 36.87 % |
| | Age (months) | -0.59 | [-1.95, 0.72] | -0.54, 0.54 | 43.50 % |

| | | | | | |
|---|---|---|---|---|---|
| | Gender | 0.10 | [-2.38, 2.70] | -0.54, 0.54 | 30.32 % |
| **GEC** | FO State 1 | -0.30 | [-3.48, 3.07] | -0.96, 0.96 | 41.48 % |
| | FO State 2 | 3.27 | [-0.82, 7.21] | -0.96, 0.96 | 14.20 % |
| | FO State 3 | -1.16 | [-4.73, 2.66] | -0.96, 0.96 | 34.15 % |
| | FO State 4 | 1.83 | [-2.92, 6.72] | -0.96, 0.96 | 27.16 % |
| | FO State 5 | -1.10 | [-6.45, 3.95] | -0.96, 0.96 | 28.70 % |
| | FO State 6 | -0.88 | [-3.73, 1.82] | -0.96, 0.96 | 42.94 % |
| | Age (months) | 0.23 | [-2.02, 2.37] | -0.96, 0.96 | 57.53 % |
| | Gender | 0.47 | [-3.21, 4.30] | -0.96, 0.96 | 36.94 % |
| **ERC** | FO State 1 | -0.77 | [-3.55, 2.07] | -0.66, 0.66 | 30.53 % |
| | FO State 2 | -0.50 | [-3.60, 2.48] | -0.66, 0.66 | 30.34 % |
| | FO State 3 | 1.46 | [-1.78, 4.65] | -0.66, 0.66 | 23.72 % |
| | FO State 4 | -0.33 | [-4.65, 3.61] | -0.66, 0.66 | 24.03 % |
| | FO State 5 | -0.22 | [-4.80, 4.54] | -0.66, 0.66 | 21.89 % |
| | FO State 6 | 0.21 | [-1.95, 2.51] | -0.66, 0.66 | 40.89 % |
| | Age (months) | -0.70 | [-2.35, 0.99] | -0.66, 0.66 | 43.70 % |
| | Gender | -2.48 | [-5.82, 0.96] | -0.66, 0.66 | 14.96 % |
| **IUS** | FO State 1 | 0.19 | [-2.84, 3.16] | -0.73, 0.73 | 33.98 % |
| | FO State 2 | -0.45 | [-3.68, 2.86] | -0.73, 0.73 | 32.10 % |
| | FO State 3 | -0.88 | [-4.30, 2.53] | -0.73, 0.73 | 28.47 % |
| | FO State 4 | -0.97 | [-5.45, 3.35] | -0.73, 0.73 | 24.30 % |
| | FO State 5 | 0.58 | [-4.29, 5.22] | -0.73, 0.73 | 23.60 % |
| | FO State 6 | 1.04 | [-1.46, 3.51] | -0.73, 0.73 | 33.33 % |
| | Age (months) | -0.87 | [-2.74, 1.03] | -0.73, 0.73 | 41.08 % |
| | Gender | 0.31 | [-3.18, 3.88] | -0.73, 0.73 | 30.30 % |

**Transition probabilities (toward each state) and questionnaires**

*(CPRS, GEC, ERC, IUS) ~ Pr transition to State1 + Pr transition to State2 + … + Pr transition to State6 + age (months) + gender*

| **Response variable** | **Parameter** | **Coefficient** | **89% HDI** | **ROPE range** | **% ROPE** |
|---|---|---|---|---|---|
| **CPRS** | Pr transition to State 1 | 0.33 | [-1.69, 2.29] | -0.54, 0.54 | 36.94 % |
| | Pr transition to State 2 | 3.82 | [1.51, 6.12] | -0.54, 0.54 | 0.00 % |
| | Pr transition to State 3 | -0.67 | [-2.66, 1.29] | -0.54, 0.54 | 33.04 % |
| | Pr transition to State 4 | 0.91 | [-0.81, 2.62] | -0.54, 0.54 | 30.56 % |
| | Pr transition to State 5 | -0.50 | [-2.15, 1.19] | -0.54, 0.54 | 39.77 % |
| | Pr transition to State 6 | -2.07 | [-3.68, -0.43] | -0.54, 0.54 | 1.51 % |
| | Age (months) | -0.42 | [-1.65, 0.93] | -0.54, 0.54 | 50.35 % |
| | Gender | -0.38 | [-2.77, 1.96] | -0.54, 0.54 | 31.57 % |
| **GEC** | Pr transition to State 1 | -0.77 | [-3.84, 2.23] | -0.96, 0.96 | 41.56 % |
| | Pr transition to State 2 | 5.78 | [1.52, 9.76] | -0.96, 0.96 | 0.00 % |
| | Pr transition to State 3 | -1.72 | [-5.08, 1.48] | -0.96, 0.96 | 30.43 % |
| | Pr transition to State 4 | 0.66 | [-2.16, 3.32] | -0.96, 0.96 | 45.04 % |
| | Pr transition to State 5 | -0.30 | [-2.87, 2.34] | -0.96, 0.96 | 50.02 % |
| | Pr transition to State 6 | -2.91 | [-5.73, -0.01] | -0.96, 0.96 | 8.65 % |
| | Age (months) | 0.09 | [-2.02, 2.31] | -0.96, 0.96 | 59.19 % |
| | Gender | -0.51 | [-4.13, 3.01] | -0.96, 0.96 | 37.70 % |
| **ERC** | Pr transition to State 1 | -0.37 | [-2.78, 1.83] | -0.66, 0.66 | 38.75 % |
| | Pr transition to State 2 | -3.29 | [-5.96, -0.65] | -0.66, 0.66 | 0.18 % |
| | Pr transition to State 3 | 3.01 | [0.53, 5.55] | -0.66, 0.66 | 1.29 % |
| | Pr transition to State 4 | 0.09 | [-1.91, 1.98] | -0.66, 0.66 | 46.65 % |
| | Pr transition to State 5 | 0.03 | [-1.97, 1.96] | -0.66, 0.66 | 46.53 % |
| | Pr transition to State 6 | 2.83 | [0.98, 4.81] | -0.66, 0.66 | 0.00 % |
| | Age (months) | -0.41 | [1.88, 1.14] | -0.66, 0.66 | 53.89 % |
| | Gender | -1.68 | [-4.55, 1.13] | -0.66, 0.66 | 21.43 % |
| **IUS** | Pr transition to State 1 | 2.56 | [-0.34, 5.63] | -0.73, 0.73 | 11.51 % |
| | Pr transition to State 2 | 1.07 | [-1.81, 4.14] | -0.73, 0.73 | 29.99 % |
| | Pr transition to State 3 | -2.67 | [-5.70, 0.33] | -0.73, 0.73 | 10.62 % |

| | | | | | |
|---|---|---|---|---|---|
| | Pr transition to State 4 | 0.97 | [-1.46, 3.39] | -0.73, 0.73 | 35.18 % |
| | Pr transition to State 5 | -0.63 | [-2.99, 1.85] | -0.73, 0.73 | 38.44 % |
| | Pr transition to State 6 | -0.10 | [-2.32, 2.16] | -0.73, 0.73 | 44.79 % |
| | Age (months) | -0.64 | [-2.59, 1.28] | -0.73, 0.73 | 45.31 % |
| | Gender | 0.44 | [-2.91, 3.68] | -0.73, 0.73 | 31.65 % |

#### S6 - Individual neural variability and neuropsychological measures

##### Switching rates and neuropsychological measures

*(fluency, Wrap time, CPM) ~ SR + age (months) + gender*

| Response variable | Parameter | Coefficient | 89% HDI | ROPE range | % ROPE |
|---|---|---|---|---|---|
| **Phonological fluency** | SR | -0.34 | [-0.75, 0.04] | -0.15, 0.15 | 17.03 % |
| | Age (months) | 0.52 | [0.14, 0.90] | -0.15, 0.15 | 0.53 % |
| | Gender | 0.13 | [-0.63, 0.92] | -0.15, 0.15 | 26.70 % |
| **Wrap time** | SR | 0.98 | [-2.46, 4.43] | -1.80, 1.80 | 5.93 % |
| | Age (months) | -1.10 | [-4.69, 2.32] | -1.80, 1.80 | 5.65 % |
| | Gender | -0.79 | [-5.71, 3.82] | -1.80, 1.80 | 4.63 % |
| **CPM** | SR | -0.01 | [-1.16, 1.09] | -0.50, 0.50 | 18.75 % |
| | Age (months) | 2.88 | [1.79, 4.06] | -0.50, 0.50 | 0.00 % |
| | Gender | -0.83 | [-2.85, 1.24] | -0.50, 0.50 | 8.35 % |

##### Entropy rates and neuropsychological measures

*(fluency, Wrap time, CPM) ~ entropy + age (months) + gender*

| Response variable | Parameter | Coefficient | 89% HDI | ROPE range | % ROPE |
|---|---|---|---|---|---|
| **Phonological fluency** | Entropy | -0.36 | [-0.76, 0.02] | -0.15, 0.15 | 15.17 % |

| | | | | | |
|---|---|---|---|---|---|
| | Age (months) | 0.52 | [0.14, 0.89] | -0.15, 0.15 | 0.26 % |
| | Gender | 0.16 | [-0.61, 0.92] | -0.15, 0.15 | 26.22 % |
| **Wrap time** | Entropy | 0.92 | [-2.62, 4.44] | -1.80, 1.80 | 63.28 % |
| | Age (months) | -1.10 | [-4.50, 2.37] | -1.80, 1.80 | 62.37 % |
| | Gender | -0.83 | [-5.73, 4.31] | -1.80, 1.80 | 53.01 % |
| **CPM** | Entropy | 0.03 | [-1.06, 1.19] | -0.50, 0.50 | 58.61 % |
| | Age (months) | 2.89 | [1.77, 3.97] | -0.50, 0.50 | 0.00 % |
| | Gender | -0.84 | [-2.89, 1.25] | -0.50, 0.50 | 28.30 % |

**Fractional occupancies and neuropsychological measures**

*(fluency, Wrap time, CPM) ~ FO state1 + FO state2 + … + FO state6 + age (months) + gender*

| Response variable | Parameter | Coefficient | 89% HDI | ROPE range | % ROPE |
|---|---|---|---|---|---|
| **Phonological fluency** | FO State 1 | 0.25 | [-1.21, 1.82] | -0.15, 0.15 | 13.67 % |
| | FO State 2 | -0.24 | [-2.15, 1.70] | -0.15, 0.15 | 11.15 % |
| | FO State 3 | -0.12 | [-1.85, 1.69] | -0.15, 0.15 | 12.35 % |
| | FO State 4 | 0.59 | [-1.54, 2.84] | -0.15, 0.15 | 9.12 % |
| | FO State 5 | -0.17 | [-4.11, 3.79] | -0.15, 0.15 | 5.74 % |
| | FO State 6 | 0.24 | [-0.82, 1.23] | -0.15, 0.15 | 19.64 % |
| | Age (months) | 0.50 | [0.08, 0.92] | -0.15, 0.15 | 3.53 % |
| | Gender | 0.16 | [-0.77, 1.03] | -0.15, 0.15 | 22.90 % |
| **Wrap time** | FO State 1 | 1.43 | [-3.07, 6.04] | -1.80, 1.80 | 52.09 % |
| | FO State 2 | 0.09 | [-4.35, 4.65] | -1.80, 1.80 | 57.57 % |
| | FO State 3 | 1.25 | [-3.26, 6.04] | -1.80, 1.80 | 52.84 % |
| | FO State 4 | 0.53 | [-4.50, 5.97] | -1.80, 1.80 | 51.87 % |
| | FO State 5 | -0.76 | [-5.88, 4.65] | -1.80, 1.80 | 51.53 % |
| | FO State 6 | -2.02 | [-6.18, 2.18] | -1.80, 1.80 | 48.75 % |

| | | | | | |
|---|---|---|---|---|---|
| | Age (months) | -1.08 | [-4.59, 2.38] | -1.80, 1.80 | 62.47 % |
| | Gender | -0.83 | [-5.96, 4.27] | -1.80, 1.80 | 51.06 % |
| **CPM** | FO State 1 | -0.76 | [-3.27, 1.69] | -0.50, 0.50 | 25.69 % |
| | FO State 2 | -0.11 | [-2.72, 2.48] | -0.50, 0.50 | 28.06 % |
| | FO State 3 | 0.18 | [-2.45, 2.69] | -0.50, 0.50 | 28.33 % |
| | FO State 4 | -1.82 | [-5.70, 2.35] | -0.50, 0.50 | 15.26 % |
| | FO State 5 | 0.90 | [-3.74, 5.32] | -0.50, 0.50 | 16.67 % |
| | FO State 6 | -0.28 | [-2.19, 1.40] | -0.50, 0.50 | 37.92 % |
| | Age (months) | 2.97 | [1.75, 4.16] | -0.50, 0.50 | 0.00 % |
| | Gender | -0.90 | [-3.21, 1.49] | -0.50, 0.50 | 25.21 % |

**Transition probabilities (toward each state) and neuropsychological measures**

*(fluency, Wrap time, CPM) ~ Pr transition to State1 + Pr transition to State2 + … + Pr transition to State6 + age (months) + gender*

| Response variable | Parameter | Coefficient | 89% HDI | ROPE range | % ROPE |
|---|---|---|---|---|---|
| **Phonological fluency** | Pr transition to State 1 | -0.29 | [-1.05, 0.51] | -0.15, 0.15 | 22.10 % |
| | Pr transition to State 2 | -0.42 | [-1.13, 0.29] | -0.15, 0.15 | 18.79 % |
| | Pr transition to State 3 | -0.29 | [-1.01, 0.39] | -0.15, 0.15 | 23.86 % |
| | Pr transition to State 4 | -0.45 | [-1.05, 0.14] | -0.15, 0.15 | 17.46 % |
| | Pr transition to State 5 | 0.52 | [-0.07, 1.17] | -0.15, 0.15 | 12.89 % |
| | Pr transition to State 6 | 0.41 | [-0.09, 0.91] | -0.15, 0.15 | 16.49 % |
| | Age (months) | 0.51 | [0.08, 0.94] | -0.15, 0.15 | 3.88 % |
| | Gender | 0.31 | [-0.57, 1.17] | -0.15, 0.15 | 20.76 % |
| **Wrap time** | Pr transition to State 1 | 0.56 | [-3.77, 4.93] | -1.80, 1.80 | 57.60 % |
| | Pr transition to State 2 | -1.40 | [-5.83, 2.95] | -1.80, 1.80 | 53.14 % |
| | Pr transition to State 3 | 1.05 | [-3.09, 5.25] | -1.80, 1.80 | 56.75 % |
| | Pr transition to State 4 | -1.03 | [-5.21, 3.06] | -1.80, 1.80 | 57.49 % |

| | | | | | |
|---|---|---|---|---|---|
| | Pr transition to State 5 | -2.44 | [-6.52, 1.92] | -1.80, 1.80 | 42.20 % |
| | Pr transition to State 6 | -0.96 | [-4.90, 2.86] | -1.80, 1.80 | 59.89 % |
| | Age (months) | -1.55 | [-5.05, 2.33] | -1.80, 1.80 | 54.96 % |
| | Gender | -1.02 | [-5.99, 4.13] | -1.80, 1.80 | 52.25 % |
| **CPM** | Pr transition to State 1 | -1.66 | [-3.67, 0.41] | -0.50, 0.50 | 13.92 % |
| | Pr transition to State 2 | -0.34 | [-2.22, 1.58] | -0.50, 0.50 | 35.80 % |
| | Pr transition to State 3 | 1.05 | [-0.81, 2.93] | -0.50, 0.50 | 25.22 % |
| | Pr transition to State 4 | -1.09 | [-2.78, 0.54] | -0.50, 0.50 | 25.68 % |
| | Pr transition to State 5 | 1.02 | [-0.61, 2.73] | -0.50, 0.50 | 26.48 % |
| | Pr transition to State 6 | 0.28 | [-1.15, 1.68] | -0.50, 0.50 | 46.12 % |
| | Age (months) | 2.79 | [1.54, 4.03] | -0.50, 0.50 | 0.00 % |
| | Gender | -0.35 | [-2.63, 1.86] | -0.50, 0.50 | 31.06 % |
